# Adaptive Ensemble Refinement of Protein Structures in High Resolution Electron Microscopy Density Maps with Radical Augmented Molecular Dynamics Flexible Fitting

**DOI:** 10.1101/2021.12.07.471672

**Authors:** Daipayan Sarkar, Hyungro Lee, John W. Vant, Matteo Turilli, Josh V. Vermaas, Shantenu Jha, Abhishek Singharoy

**Affiliations:** MSU-DOE Plant Research Laboratory, East Lansing, MI, USA 48824; School of Molecular Sciences, Arizona State University, Tempe, AZ, USA 85281; Pacific Northwest National Laboratory, Richland, WA, USA 99354; Rutgers University, Electrical & Computer Engineering, New Brunswick, NJ, USA 08854; Brookhaven National Laboratory, Computational Science Initiative, Upton, NY, USA 11973

## Abstract

Recent advances in cryo-electron microscopy (cryo-EM) have enabled modeling macromolecular complexes that are essential components of the cellular machinery. The density maps derived from cryo-EM experiments are often integrated with manual, knowledge or artificial intelligence driven, and physics-guided computational methods to build, fit, and refine molecular structures. Going beyond a single stationary-structure determination scheme, it is becoming more common to interpret the experimental data with an ensemble of models, which contributes to an average observation. Hence, there is a need to decide on the quality of an ensemble of protein structures on-the-fly, while refining them against the density maps. We introduce such an adaptive decision making scheme during the molecular dynamics flexible fitting (MDFF) of biomolecules. Using RADICAL-Cybertools, and the new RADICAL augmented MDFF implementation (R-MDFF) is examined in high-performance computing environments for refinement of two protein systems, Adenylate Kinase and Carbon Monoxide Dehydrogenase. For the test cases, use of multiple replicas in flexible fitting with adaptive decision making in R-MDFF improves the overall correlation to the density by 40% relative to the refinements of the brute-force MDFF. The improvements are particularly significant at high, 2 - 3 Å map resolutions. More importantly, the ensemble model captures key features of biologically relevant molecular dynamics that is inaccessible to a single-model interpretation. Finally, the pipeline is applicable to systems of growing sizes, which is demonstrated using ensemble refinement of capsid proteins from Chimpanzee adenovirus. The overhead for decision making remaining low and robust to computing environments. The software is publicly available on GitHub and includes a short user guide to install the R-MDFF on different computing environments, from local Linux based workstations to High Performance Computing (HPC) environments.

## 1 Introduction

Integrative modeling is an area of rapid methodological developments, wherein, atom-resolved structure(s) of biological systems are determined by merging data from multiple experimental sources with physics ^1–3^ and informatics-guided approaches. ^4^ These elegant fitting, ^1–3,5–10^ learning ^11^ sand inferencing ^12–16^ methodologies have been successful in resolving a range of structures, starting with soluble and membrane proteins up to sub-cellular complex architectures. ^17,18^ Integrative models routinely make it to top positions at the EMDB and PDB competitions, serving a diverse cross-section of the Biophysics community. ^19^ Advances in protein structure modeling, ^20–23^ evolutionary covariance or multi-sequence alignments offer excellent constraints for initiating such hybrid pipelines. ^24^

A key issue in integrating structural or biochemical information with simulations stems from the heterogeneity of the data. The data quality can be spatially variant, spanning anywhere between coarse-grained to near-atomistic level of details. As a natural consequence of this heterogeneity, a single-model interpretation of the experimental data becomes implausible, opening the door to an ensemble treatment of the data. ^25^ The ensemble models capture on one hand, the most probable interpretation of the data, while on the other, pinpoints rare-events and hidden conformations. Biology often employs such conformational diversity in problems of allostery and recognition, motivating the refinement of experimental knowledge against molecular ensembles. ^14^ While deep learning tools have started offering high-quality protein structural predictions, it is also demonstrated that the per-residue uncertainty metrics determined from such predictions provide useful prior information for the generation of conformational ensembles. ^14,26,27^ Among structural determination experiments, cryo-EM data has inspired such ensemble interpretation for some years now ^18,28^ leading to the development data-guided entropy maximization pipelines ^14,26,27^ and conformational search pipelines. ^6,29–31^ Another advantage of the ensemble interpretation is that, the generation of multiple independent atomic models using an EM density and statistical analysis of their map-model agreement offers metrics of global as well as local EM map quality. ^32^ This ensemble approach offers essentially both a quantitative and qualitative assessment of the precision of the models and their representation of the density. However, integration of routinely observed conformationally heterogeneous cryo-EM data and molecular simulations require a number of adaptive decision-making steps, challenging their massively parallel implementation. Here, a computing workflow is presented that can be generally leveraged for hybrid modeling endeavors.

The size of the ensembles that collectively describes the diversity in single-particle images is representative of conformational and compositional heterogeneity.^29^ For proteins of molecular mass 500 kDa or bigger, composed of 5000 residues or more, a single CPU is expected to take 5000 years of wall-clock time for sampling the conformational ensembles using either brute force molecular dynamics (MD) or Monte Carlo simulations; ^33^ even the fastest GPUs of the day will not rescue this situation. Alternatively, data-guided enhanced sampling methodologies, such as MELD ^13^ (integrated with NAMD via the recently completed CryoFold plugin ^14^) or backbone tracing methodologies such as MAINMAST^34^ and analogous methods, ^11^ by themselves, either remain system-size limited, generating ensembles for only local regions within a map, or require further refinements using conjugate gradient minimization or free-energy schemes ^35^ to determine thermodynamic ensemble. As a step towards addressing this issue, by leveraging classical force fields (so-called CHARMM ^36^ energy functions) we have developed a range of molecular dynamics flexible fitting (MDFF) methodologies for integrating x-ray and cryo-EM data with MD simulations. ^1–3,37^ The simulations are biased towards conforming molecular models into forms consistent with the experimental density maps.

These protocols are available through MD simulation engine NAMD, ^38^ recently to GROMACS ^6^ and are also expanded as plugins, such as ISOLDE ^39^ in ChimeraX. ^40,41^ As a natural outcome of this fitting procedure, the most probable data-guided models are derived, e.g. for complex systems like the ribosomal machinery, virus particles and membrane proteins. ^18^ However, the conformational heterogeneity^29,42,43^ that contributes to the uncertainty of the experimental data is lost.

In this article we explore whether, it is possible to recover portions of the conformations lost in brute-force MDFF by running multiple replicas of MDFF in parallel with ‘adaptive decision making’. Notably, adaptive sampling is one of the key methods used for enhancing MD simulations to model free energy changes. ^44,45^ Rather than physically enforcing a model into a map, this approach skews the probability of an ensemble of models towards maximizing their consistency with the map. This way, there remains a finite probability of visiting several uncertain structures, while still emphasizing determination of the most probable molecular models.

Traditional high-performance computing (HPC) approaches fail to make data-driven decisions within a multi-replica ensemble modeling workflow. We employ RADICAL-Cybertools, ^46^ and in particular Ensemble Toolkit (EnTK), ^47^ to overcome this challenge of developing multi-replica MDFF as a workflow application. Herein, EnTK deploys an application programming interface (API) for casting the MDFF simulation and analysis workflow as a hierarchy of pipelines, stages and tasks. Simultaneously, the RADICAL-Pilot (RP) ^48,49^ is employed as the high-performance and dynamic resource management layer. This workflow identifies all the flexible fitting tasks within a pipeline, acquires heterogeneously distributed resources to complete multiple parallel pipelines, and manages the overall execution of the stages iteratively.

A classical approach involves long brute-force MD simulations that are often stuck in local minima, requiring additional steps to find interesting regions of the conformational search space. In contrast, adaptive sampling implements an iterative loop that concurrently executes multiple simulations, each with short simulation time. ^50–52^ Goodness-of-fit metrics map/model are analyzed at every iteration - this analysis increases the probability of finding models that are consistent with the data, reducing the possibilities of getting trapped in any local energy minimum. The decision to enhance the sampling of specific models can be based a number of map-model metrics, ^19^ such as TM scores, ^53^ MolProbity, ^54^ EMRinger, ^55^ Q-score. ^56^ In our first proof-of-concept adaptive MDFF workflow, a simple global cross correlation coefficient (CC) is employed as a criteria to guide the choice of refinement models; Molprobilty statistics are employed for cross-validation.

Outlined in **Figure 1**, workflow application using R-MDFF composes individual simulations and supports analysis calculation on intermediate results to perform adaptive sampling. The scheme iteratively screens model populations based on their CCs with the map, and improves efficiency of computing resource consumption over longer simulations. We find that, powered by EnTK’s data-staging capabilities and check-pointing of the parallel MD simulations available on NAMD, MDFF trajectories intermittently screened by CC values offer an ensemble of refined models. We have tested the adaptive MDFF workflow with up to 100 replicas, each encompassing 16 iterations across resolutions of 2 Å to 5 Å, and achieved around significant improvement in map-model fitting over a long brute-force simulation. Similar efficiencies are noted while comparing the adaptive workflow with multi-replica, yet non-adaptive implementations of MDFF. The pipeline is further tested for using up to 400 replica with (1 node/replica). In all these cases, we find that an ensemble approach with adaptive decision making offers more diverse ensembles than brute force MDFF. Thus, going beyond traditional MDFF, these ensembles capture on one hand, the ‘best’ model, while simultaneously the uncertainty in the assignments on the other. Remarkably, the performance of the workflow improves with system-size (3341 atoms in Adenylate Kinase and 11452 atoms in Carbon Monoxide Dehydrogenase atoms), where enhanced sampling finds more low-probability structures than brute-force MD and efficiency remains robust to computing platforms. Ensemble refinements are further extended to study poorly resolved components of Chimpanzee adenovirus ChAdOx1. Taken together, our implementation breaks free of the traditional high-performance computing execution model that assumes singular jobs and static execution of tasks and data, to one that is fundamentally designed for data-integration and assimilation across different scales, quality and sparsity. The cryo-EM community has actively sought ways of extracting not just stationary structures, but ensembles and more importantly, molecular dynamics information from electron density data are ever-increasing. ^29,30,57,58^

**Figure 1:**
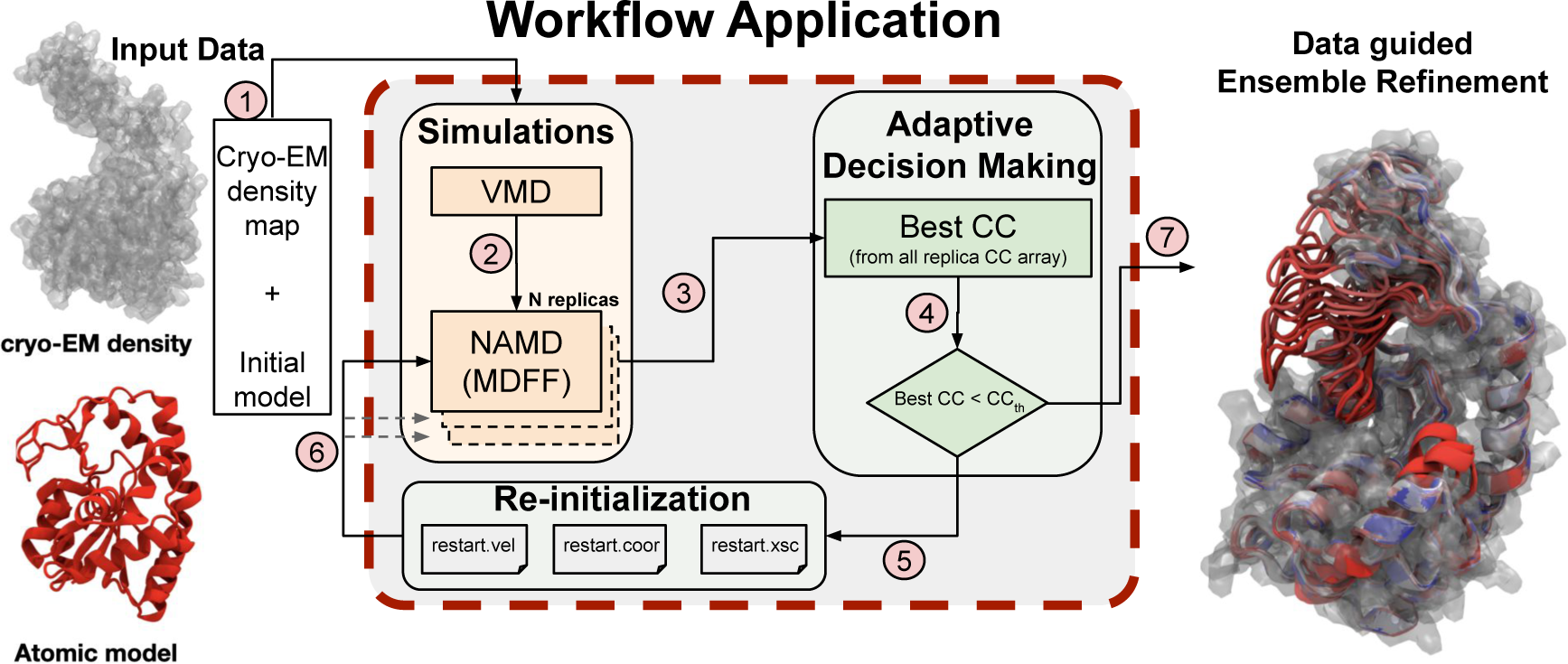
Overview of the workflow application for R-MDFF. The schematic shows how NAMD/VMD is used to perform flexible fitting iteratively. Internal boxes with annotation numbers indicate the sequence of the workflow: (1) Input data (2) Simulation preparation and execution using VMD and NAMD respectively (3) Building CC matrix and sort through CC matrix to select best CC (4) Check if best CC is lower than threshold CC (5) Use the current state of the molecular sytem corresponding to the best CC using the restart files (6) Re-seed all replicas with the restart files and perform the next iteration of flexible fitting (7) Data guided ensemble refined models.

## 2 Adaptive Integrative Modeling using R-MDFF

Adaptive decision making for the refinement of multiple protein structures coupled to 3-dimensional (3D) electron density maps is implemented as an iterative simulation-analysis workflow enabled by EnTK (**Figure 1** and **Algorithm 1**). Within R-MDFF, a workflow is defined as an ensemble of *simulation* + *analysis* pipelines that synchronously execute on HPC resources. Each constituent pipeline enable seven serial tasks: **(1)** load an empirically determined density map or generate a simulated map. Then convert this map to an MDFF potential, Eq. 1 and 2. Independently, examine the quality of an initial search model in terms of stereochemical properties, and perform rigid body docking to place this search model inside the EM density map, e.g. using Chimera ^41^ or Situs. ^59^ **(2)** Define the secondary structure restraints. Visual Molecular Dynamics (VMD) ^60^ then prepares the input files required by NAMD ^38,61^ to deploy MDFF. Multiple replicas of the system are prepared, under the same R-MDFF pipeline, as shown in **Figure 1**. Then, the ensemble of MDFF simulations are performed in parallel. **(3)** VMD’s scripting interface is re-used to calculate the interim cross-correlation value between the atomic models from each replica and the EM density map. The CC values are extracted from VMD log files for different replicas are then combined to construct a matrix. EnTK uses a data staging area to move this matrix from flexible fitting to the adaptive decision-making block across multiple replicas. **(4)** Here, a decision is made on whether the flexible fitting simulations will continue or terminate, based on whether the computed CC is greater than or equal to the user-defined threshold. This on-the-fly map-model analysis enables an adaptive flexible fitting algorithm to run recursively inside EnTK, without user intervention. When the threshold CC is not met at the end of tasks 1-4, subsequent iterations are performed, wherein **(5)** all replicas are reseeded with the atom coordinates, velocities, and periodic system information corresponding to MDFF model with the best CC from the previous iteration, and the next round of multi-replica MDFF proceeds. If the map correlation of the best-fitted model decreases along the forward iteration, the new poorly fitted starting conformation is accepted with a weight *min* (1, *e*^Δ^*^ECC/KT^*).

Here, we follow Δ*E_CC_* = *k*(*CC_N_*_+1_ *− CC_N_*) for iteration no. *N* and *k* = 5 *×* 10^5^ kJ/mol. ^6^ For a failed move, the fitting restarts with the initial conformations of the last iteration, and the criterion is reused to find a new starting structure. **(6)** Again, EnTK uses data staging area to store these information in files and provide them to the replicas. This feature not only makes the algorithm adaptive, but offers future scope of improvement for applications requiring advanced decision-making, either based on inferencing ^12–14,42,62^ or neural network based machine learning algorithms. ^63–67^ Finally, the application converges to yield a refined ensemble **(7)**, which exit the R-MDFF workflow and downloads results to the end-user’s working directory.

**Algorithm 1:**
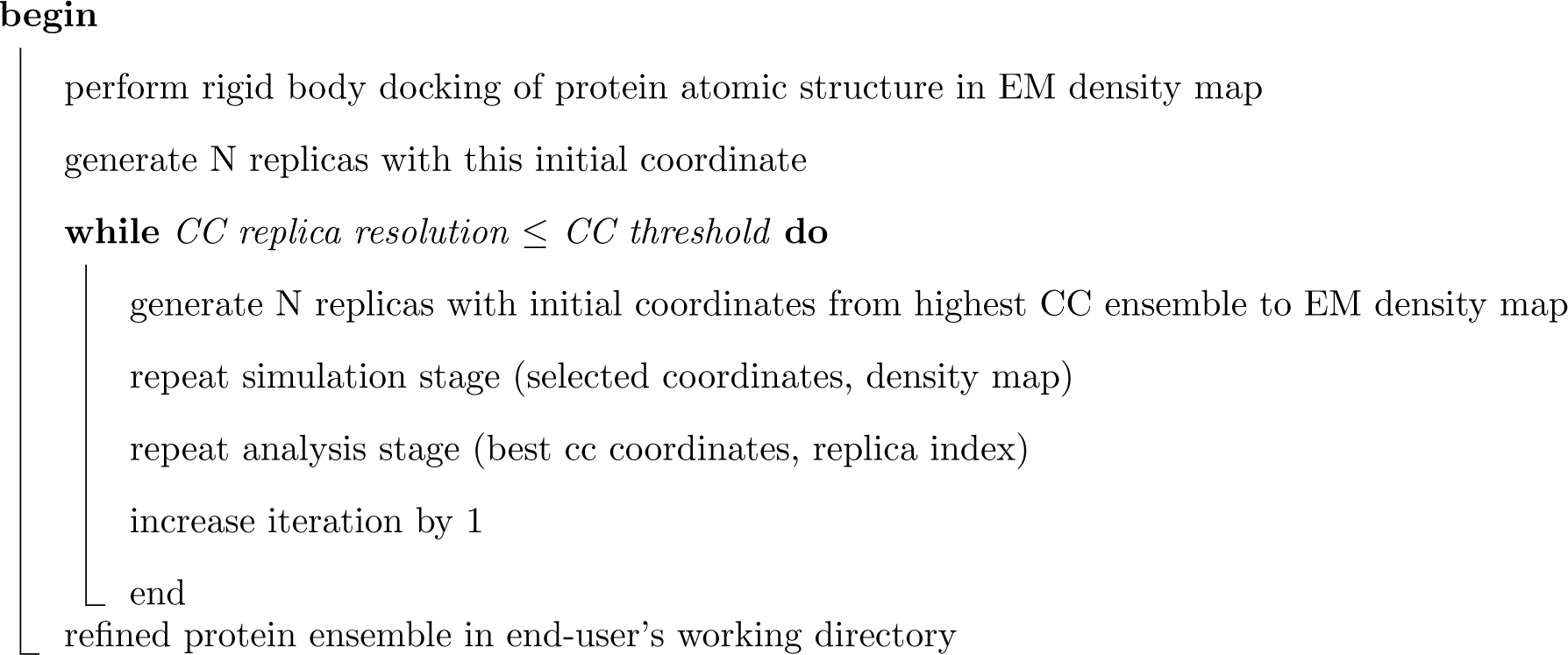
R-MDFF scheme with adaptive decision making based on CC.

The R-MDFF API is implemented as a Python module, loaded into the workflow application’s code. ^68^ The API exposes classes for pipeline, stage, and task, allowing one to directly map the workflow description to the logical representation of an ensemble of simulations. Each task object exposes a set of variables with which to configure input, output files, executable, resource requirements, and pre/post execution directives.

Finally, an application manager object is used to contain the workflow description and execute it with a single AppManager.run() method. The iteration logic to change the workflow description and issue another AppManager.run() is written in pure Python as part of the workflow application. The entire R-MDFF workflow application of this paper required only 500 lines of Python code. ^68^

As already described in Balasubramanian et al. ^47^, EnTK complements the ensemble simulation paradigm with decision making through real-time workflow and parameter changes, based on the results of the analysis stages. In the present context, this feature enables iterative workflow executions with a single HPC batch-job submission, avoiding costly manual evaluation of cross-correlation coefficient, workflow editing, and re-submission, as demonstrated here for flexibly fitting biomolecules in cryo-EM density maps. EnTK also abstracts from the users the need to explicitly manage data flow and task execution. It manages data staging so that each task of each stage has either a copy or a link to all the NAMD input files it requires, allowing the users to focus on the MDFF simulation and VMD analysis methods, without having to explicitly handle data sourcing, saving, and exchange. Furthermore, EnTK schedules and executes the workflow’s tasks, managing the mapping of tasks to available resources on each compute node allocated to the workflow execution. Users only have to specify the amount of CPU cores/GPUs needed by each task and whether the task is (Open)MPI.

## 3 Methods

Modern adaptive sampling frameworks are dynamic, extensible, scalable and robust to facilitate hundreds or thousands of experiments for searching different structures, and specialized features can be added to solve existing problems through the framework. We developed a workflow application using RADICAL-Cybertools ^46^ that provides a scalable workflow framework for implementing ensemble refinement with cross correlation calculation on HPC computing resources. R-MDFF (RADICAL-Cybertools enhanced Molecular Dynamics Flexible Fitting), depicted in **Figure 1**, supports adaptive decision making algorithms to iterate between molecular dynamics flexible fitting simulation and cross-correlation analysis. Our workflow application is portable to explore the space of experimental configurations and support various use cases, so that the ensemble refinement produces results on different dimensions of a physical system; resolution density, simulation length, replica count, and HPC resource. The complete integration is explained in the following sections: (a) MDFF simulation, (b) CC analysis, (c) RADICAL-cybertools and (d) validation approaches.

### 3.1 Molecular Dynamics Flexible Fitting simulation

In the pipeline simulation stage, R-MDFF uses the conventional MDFF algorithm, as described in. ^3^ Briefly, MDFF requires, as input data, an initial structure and a cryo-EM density map. A potential map is generated from the density and subsequently used to bias a MD simulation of the initial structure. The structure is subject to the EM-derived potential while simultaneously undergoing structural dynamics as described by the MD force field.

Let the Coulomb potential associated with the EM map be Φ(**r**). Then the MDFF potential map is given by,

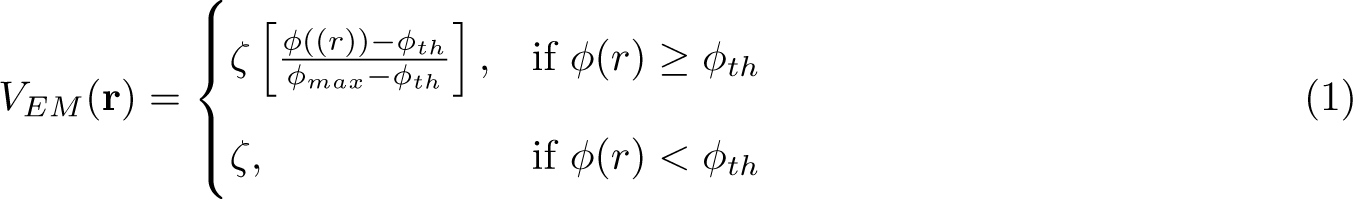

where *ζ* is a scaling factor that controls the strength of the coupling of atoms to the MDFF potential, *φ_th_*is a threshold for disregarding noise, and *φ_max_* = *max*(*φ*(*r*)). The potential energy contribution from the MDFF forces is then

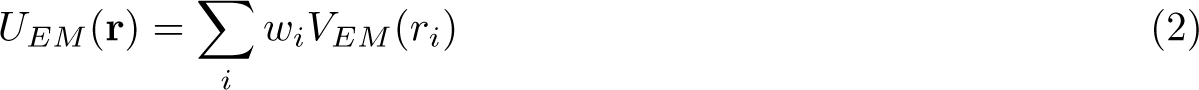

where *i* labels the atoms in the structure and *w_i_* is an atom-dependent weight, usually the atomic mass. During the simulation, the total potential acting on the system is given by,

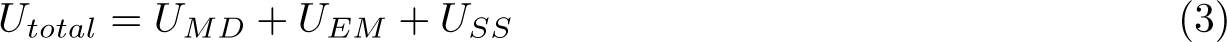

where *U_MD_*is the MD potential energy as provided by MD force fields (e.g. CHARMM) and *U_SS_* is a secondary structure restraint potential that prevents warping of the secondary structure by the potentially strong forces due to *U_EM_* . ^1,69^ A detailed description of the MDFF methodology is presented in. ^1,2^ Specific simulation parameters for the example cases of ADK and CODH are provided on the GitHub page. ^68^

### 3.2 Cross-correlation analysis

The analysis and decision-making part of the ensemble refinement involves calculating the map-model cross correlation (CC) value for all replicas at every iteration of the R-MDFF workflow. For *t* MD steps and *M* replicas per iteration, the total simulation time is equal to *t_steps/iteration_ × N_iteration_ × M_replicas_*. At the end of each iteration, the CC for *M_replicas_* number of resulting structures is computed against the target map to examine the quality of fitting. Atomic coordinates corresponding to the Monte Carlo-like selection rule described in Sect. 2 are used to restart all the replicas for the next iteration.

### 3.3 RADICAL-Cybertools

In order to implement the pipeline, we have extended an open-source, Python framework – RADICAL Ensemble Toolkit (EnTK) – that facilitates adaptive ensemble biomolecular simulations at scale. The first step of writing the EnTK workflow code is to construct a task parallel execution of MDFF simulation using NAMD 2.14, ^38,61^ and to connect the analysis stage to find highest CC values among replicas. While all the necessary information such as NAMD checkpoints and CC values are kept under EnTK’s data staging area, distributed computing resources are coordinated to ensure the workflow performance over CPUs and GPUs from heterogeneous HPC platforms. In addition, several features have been added to the application by utilizing existing capabilities of RADICAL-Cybertools. Tcl scripting is interfaced with EnTK APIs to interact with VMD software directly and a partitioned scheduling is introduced to assign a single node per replica for the best performance of NAMD simulations. Usability and productivity have been addressed to automate resource configurations and experiment settings as well as ensuring reproducibility of scientific data. The R-MDFF application integrates the NAMD engine and VMD for analysis methods and thus requires only a few lines of settings in a workflow management file without source code modifications. The application, R-MDFF is available on GitHub (https://github.com/radical-collaboration/MDFF-EnTK) and implemented to support adaptive decision making for ensemble-based simulations and to enable the novel analysis method, MDFF or others on HPC resources.

### 3.4 Other analyses - MolProbity and PCA

MolProbity ^54^ scores are calculated to determine the quality of the ensemble of structures for protein ADK after adaptive decision based flexible fitting. The distribution of MolProbity scores indicate that as the ensemble members increase, ranging from 16 to 400, a population of high quality structural ensemble (MolProbity score ∼ 0.75) is observed.

For the analysis of protein molecular dynamics simulations, principal component analysis (PCA)^70–72^ approach can monitor the individual modes, thereby allowing one to filter the major modes of collective motion from local fluctuations. Often these principal modes of motion is correlated with protein function, the reduced dimensional subspace spanned by these modes was termed essential dynamics, ^73^ reflecting the modes which correspond to essential biological function. Also, using PCA we can distinguish converged structures (fitted to EM density map) from outlier structures (outside the EM density map). ^71^

We performed PCA using ProDY^74^ and represent the essential dynamics using the Normal Mode Wiz-ard ^74^ plugin in VMD. ^60^ The principal components are calculated corresponding to 16 and 400 ensemble members.PCA results of evaluating the structural ensemble of protein ADK across multiple iterations in R-MDFF, suggests while fitting to a high resolution EM density map with low ensemble members (16 replicas), the essential dynamics capture mostly the biologically relevant ”lid-closed” state. However, increasing the number of ensemble members (400 replicas), results indicate that normal modes can probe both the the biologically relevant ”lid-open” and ”lid-closed” states.

## 4 Results and Discussion

We conducted a series of experiments using R-MDFF by varying the number of replicas and iterations, and length of flexible fitting simulations. With these parameters, we compared the quality of the refined models for two example systems, namely, adenylate kinase (ADK) and carbon monoxide dehydrogenase (CODH) proteins. The robustness of the protocol is demonstrated on two HPC facilities, namely on Oak Ridge Leadership Computing’s Summit, each node on which has two IBM Power9 processors and six NVIDIA V100 GPU accelerators, and on Pittsburgh Supercomputer Center’s Bridges2 having two AMD EPYC 7742 processors per node.

### 4.1 Adaptive decision making using R-MDFF provides variance in ensemble refinement of high-resolution density maps

We start by fitting the ‘closed’ conformation of ADK (PDB: 1AKE) into synthetic density maps derived from its ‘open’ form (PDB: 4AKE). In **Figure 2** the CC changes are presented as a function of iterations for ensemble members with 4, 8, 16 and 32 replicas, and at map resolutions of 1.8, 3 and 5 Å. Using the utility Phenix.maps and the reported structure factors, ADK density maps were constructed at the native 1.8 Å resolution, and were further truncated at intermediate (3 Å) and low (5 Å) map resolutions. The product of number of replicas (*M_replicas_*) *×* number of iterations (*N_iterations_*) *×* the flexible fitting MD steps per iteration and per replica (*t_steps/iteration/replica_*) which results in 16 nanoseconds (ns) of sampling.

**Figure 2:**
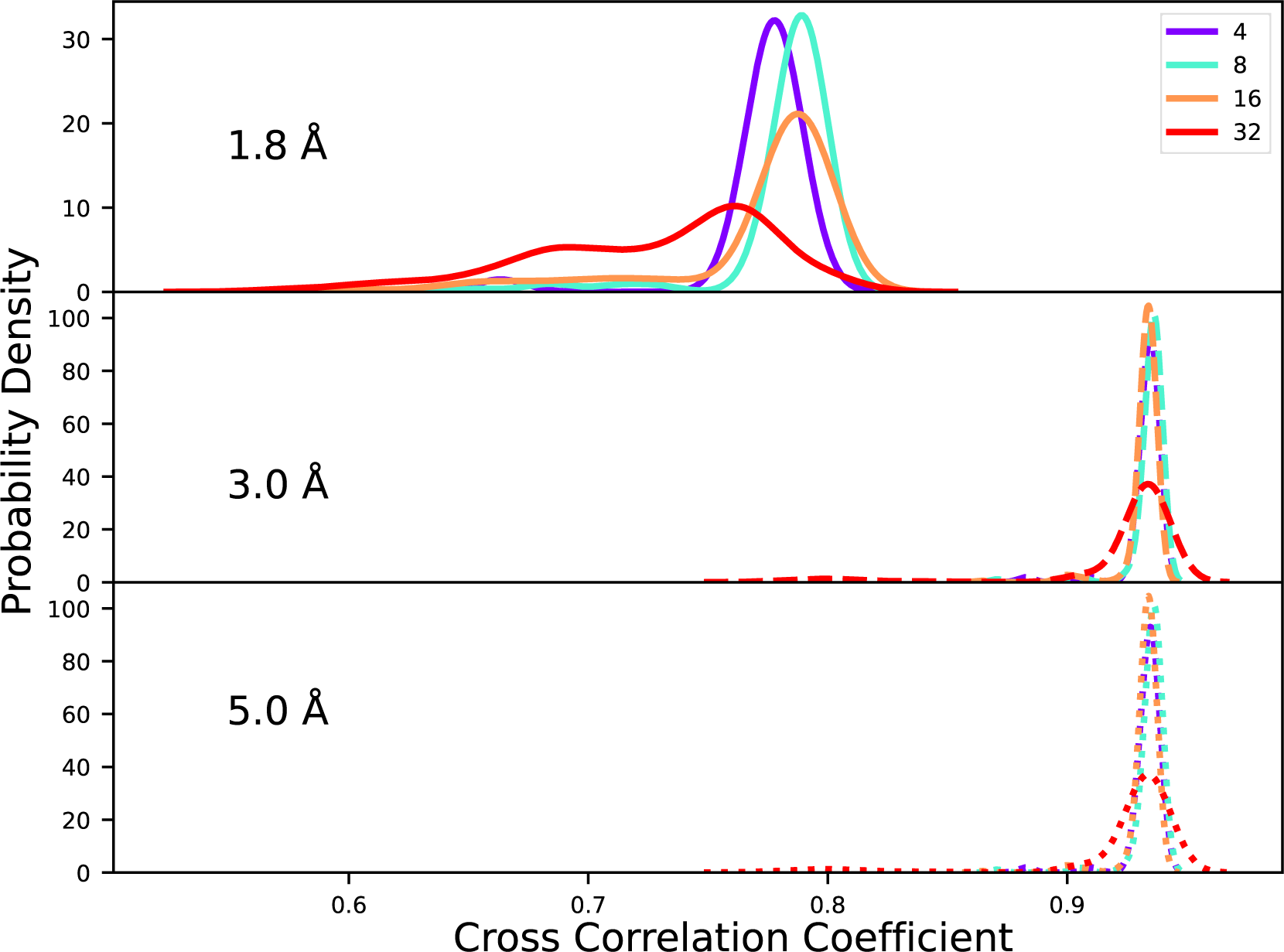
Cross correlation coefficient after flexibily fitting ADK with R-MDFF at different resolution density maps. Data presented for high resolution (1.8 Å), intermediate resolution (3.0 Å) and low resolution (5.0 Å) cryo-electron microscopy density maps, for different ensemble members, 4 (purple), 8 (aqua), 16 (orange) and 32 (red) respectively. We find that replica number directly correlates with we have larger variability amongst ensemble members for high resolution density map.

For intermediate and low resolutions, the CC peak is *≈* 0.9, while at high resolution this value shifts to *≈* 0.8. While the former took one to two iterations only, the later took up to ten iterations to reach the highest CC values. The determination of the best-fitted model is almost independent of the number of replicas used, barring at 1.8 Å where a small increase in peak CC from 0.78 to 0.80 is observed when *M_replicas_* doubled from 4 to 8. As expected, the change in the quality of fit from 0.9 (against the 3.0 Å and 5.0 Å maps) to 0.8 (against the 1.8 Å maps) stems from the many conformations of a model that can match intermediate to low resolution density, the number of which decreases at higher resolutions.

In **Figure 2**, as the replica count further increases to 16 and 32, the distribution of CC values becomes wider for a high-resolution density map. Interestingly, the best CC values at the end of every iteration does not linearly increase with iterations, but rather show fluctuations with an overall increasing trend, Figure **S1**. For example, in iterations 3 to 5 of the 32-replica refinement the CC’s decrease, suggesting that the *E_CC_*-selection criterion does not uniquely bias the distribution towards higher CC values, offering a probability to sample also the poorly fitted structures. This wider distribution implies conformational diversity ^29^ in the ensemble of refined protein structures. Conventionally, MDFF generates protein structures with low variance and high bias towards maximizing correlation with the target density. At higher resolutions, the population of the structures is further skewed. Even with the use of a relatively small number of R-MDFF replicas (*M_replicas_*= 32) encoded through EnTK, the workflow generates a range of structures with CCs between 0.78 to 0.82. On the contrary, a single long MDFF trajectory with identical cumulative time of 16 ns produces final models strongly peaked at CC ∼ 0.67. Therefore, unlike conventional flexible fitting, the R-MDFF workflow generates models that represent different propensities for large-scale protein conformations, still including the most probable model with an improvement of 22% over a single long MDFF.

### 4.2 Statistics of map-model fits improve with larger replica simulations

An improvement in ADK fit quality on a single long MDFF trajectory, particularly at 1.8 Å resolution, motivated further exploration of fit quality as a function of *M_replicas_*. First, we have repeated the *M_replicas_* = 64 computations without the adaptive decision block in R-MDFF. The most probable CC values decrease to 0.52 compared to 0.82 determined by executing the entire workflow, see **Figure 3**. This significant difference of 40% in the fit of the model to map implies that it is not just the sampling of an increased number of replicas, rather the adaptive re-initialization strategy employed every iteration successfully improves the simple MDFF results.

**Figure 3:**
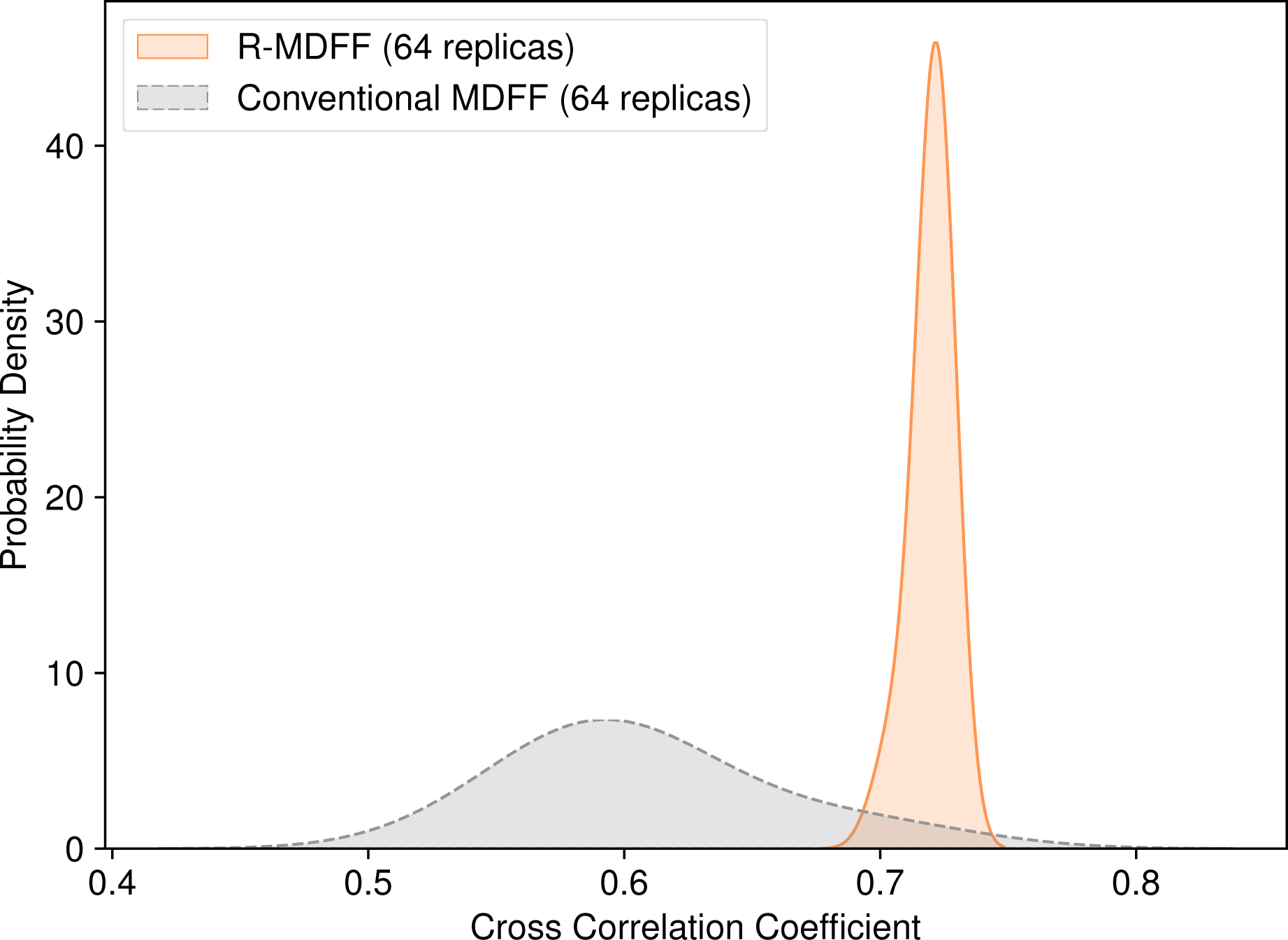
Conventional versus R-MDFF on flexbile fitting ADK in high resolution density map. Comparing cross correlation between conventional and R-MDFF when fitting ADK to high resolution EM map of 1.8 Å using 64 replicas. We find R-MDFF to perform better than a conventional MDFF, with a population of high cross correlation ensemble members, indicated by the solid line.

Next, it is examined whether increasing the simulation length per iteration for every replica, i.e. *t_steps/iteration/replica_* has an effect on improving the quality of fit for larger number of replicas. From **Figure 4**, one notice that the CC value increases and then drops and forks as the *M_replicas_* increases beyond 32. This is expected, since for higher *M_replicas_*the *t_steps/iteration/replica_*is decreased to conserve the cumulative simulation length as that of a single MDFF trajectory. To address this issue, for replicas 64, 100, 200, 400 we increase the simulation length per iteration for each replica to match that of the 16-replica workflow. We chose the *t_steps/iteration/replica_* = 16 ns from the *M_replicas_* =16 setup, as that provides some rare yet high CC peaks.

**Figure 4:**
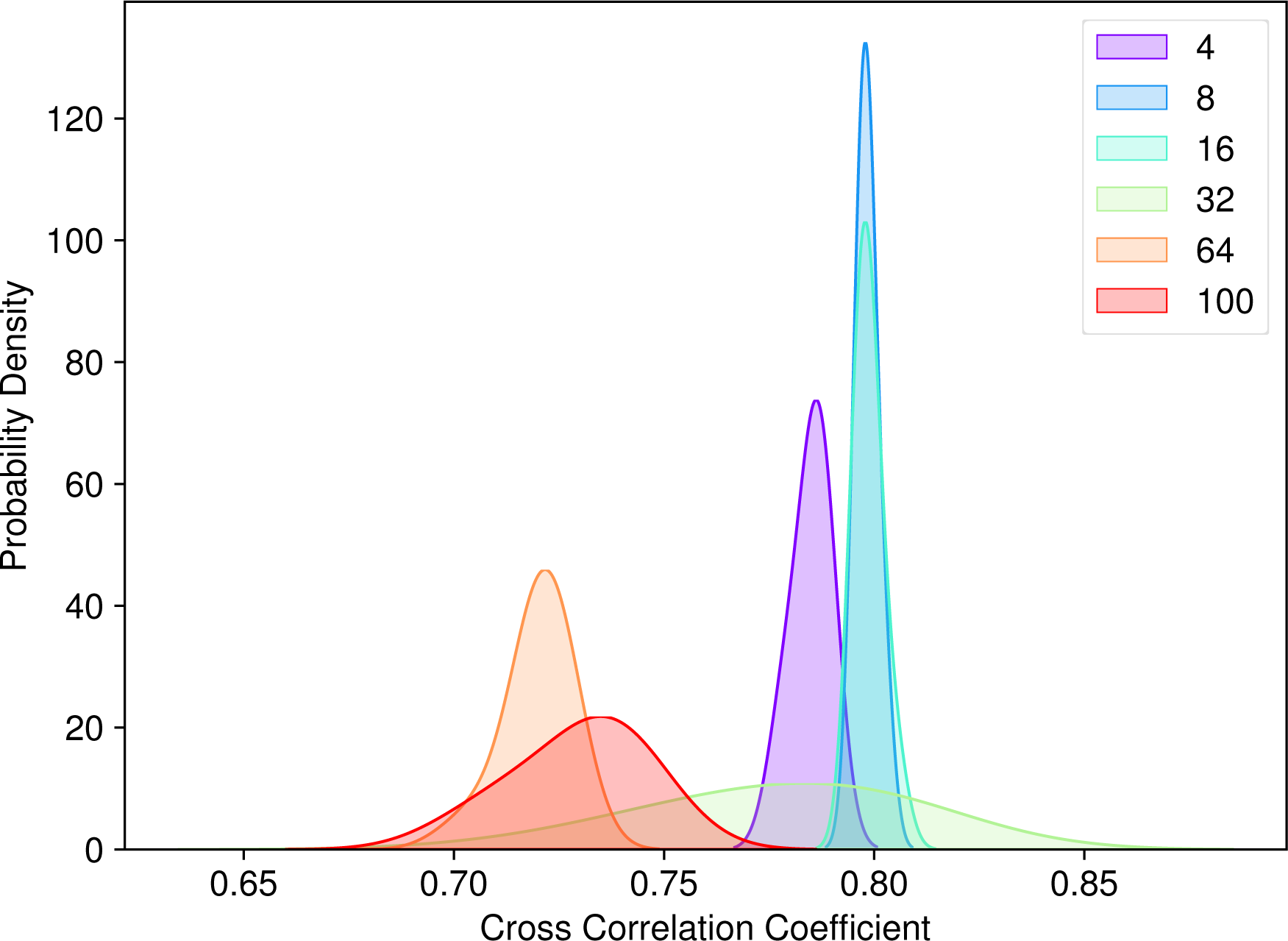
Change in quality of fitted ensemble with replica size. Best cross-correlation after iterative adaptive flexible fitting using R-MDFF, to fit ADK to a high resolution 1.8 Å map, for different ensemble sizes with *M_replicas_* = 4 - 100. Monitoring the effect of replica size on the quality of fit. We observe cross-correlation to increase till 32 replicas, after which the value starts decreasing.

As *t_steps/iteration/replica_* increased from 20 ps to 80 ps, for *M_replicas_* = 64, to match the simulation length for *M_replicas_* = 16 from **Figure 4**, the CCs improved systematically by 11% from 0.72 to 0.82 (**Figure 5**). More importantly, at the higher values of *M_replicas_*, e.g. 100, 200, and 400, a broad, and in fact, bimodal distribution of refined models is derived, while maintaining the same *t_steps/iteration/replica_* = 80 ps. The bimodal distribution captured using 400 ensemble members represents the conformational heterogeneity ^29,42,43^ observed in cryo-EM density maps. These conformational heterogeneity corresponds to different thermodynamic states of the biomolecule.

**Figure 5:**
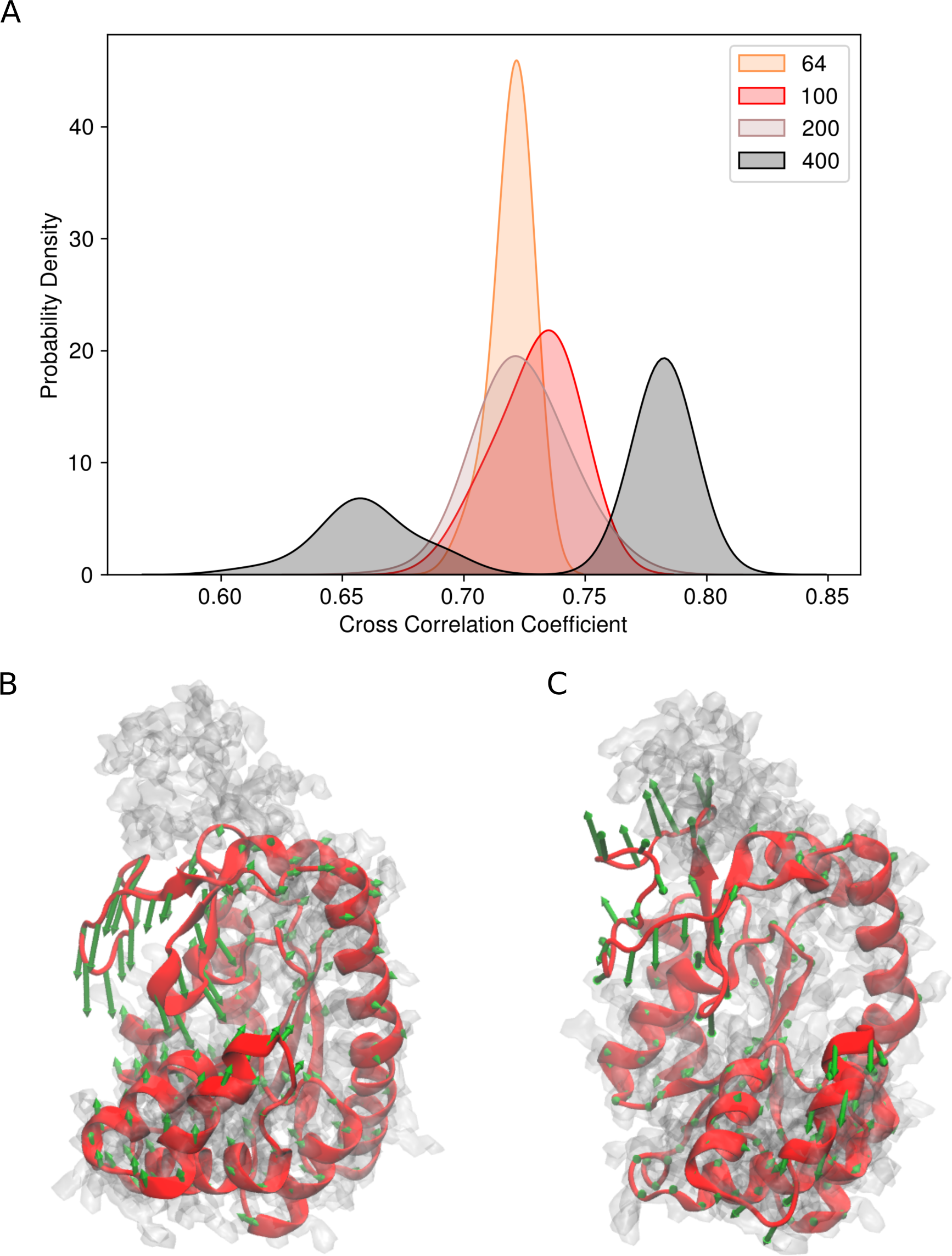
Impact of total simulation time on the quality of fit. (A) Best cross-correlation (CC) after adaptive flexible fitting of ADK to a high resolution (1.8 Å) map, for different ensemble members (64, 100, 200 and 400). (B) Principal component analysis (PCA) of ADK protein with 16 replicas flexibly fitting to 1.8 Å using adaptive protocol (C) PCA of ADK protein with 400 replicas flexibly fitting to 1.8 Å using adaptive protocol. PCA results demonstrate the experimentally observed hinge motion, can be found with a high23replica number, but not for a low replica number.

Models with high CC around 0.8 are expected given the inherent bias of the density data within MDFF simulations. These models remain within an RMSD of 1 Å relative to structures of comparable CC derived from the 16-replica simulations, see SI, **Figure S2**. However, the distribution of models isolated with statistical significance, and lower yet still decent CC values between 0.7 to 0.8, were obscured by single, smaller-replica or non R-MDFF jobs. The quality of these structures was determined employing a MolProbity ^54^ analysis of all members of the generated protein ensemble. Despite a broad distribution in the quality of fit, the quality of model remains universally high as seen through MolProbity scores peaked between 0.75 - 1.75, see SI figures, **Figures S3 - S7**. Thus, the multi-model description inferred from the 1.8 Å ADK density map remain energetically viable conformations of the protein that remain in the vicinity of the best-fitted model, but with variations that can reflect the dynamics of ADK.

In the open state of ADK, its so-called “lid” domain undergoes a hinge-like movement^35,75–77^ to maintain a conformational pre-equilibrium with the closed state, with the open state being more prevalent for apo protein. ^78^ Such movements are confirmed by transition path sampling simulations and FRET experiments. ^79,80^ Illustrated in **Figure 5**, principal components from the ensemble of converged R-MDFF models collated across 400 replicas clearly captures this hinge movement of “lid” opening and closing essential towards the biological function catalyzes the interconversion of the various adenosine phosphates (ATP, ADP, and AMP). PCAs from the ensemble peaked at 0.65 offers signatures of closing, while those peaked at 0.80 have conformed to a new stable state showing minimal chances of closing. We did not find any strong overlap between the CC=0.65 and 0.8 peaks in **Figure 5**, implying that if the initial models are too close to the fully open state it will not be able to sample the intermediate state. With further simulations, the sampling of models with CC values around 0.72 can be improved to fully see these open to less-open transitions. In contrast, a skewed distribution of the just the best-fitted models is obtained with *M_replicas_* = 16, indicating only the “lid” open state. Thus, by using a probabilistic selection criterion within R-MDFF, rather than a deterministic one used conventionally with data-guide simulations such as MDFF, the space of CCs is more exhaustively sampled during flexible fitting, and the evolution of an ensemble can be monitored to gain insights on the structure-function relationship for biomolecules.

### 4.3 Refinement protocol is robust to system size

Robustness of the newly implemented R-MDFF parameters estimated from our multi-replica ADK simulations is tested using a second example of larger protein, namely carbon monoxide dehydrogenase. The closed conformation (PDB - 1OAO: chain C) was used as the search model, while the open state (1OAO: chain D) was the target. Similar to ADK, the fitting was performed with maps of the reported 1.9 Å and synthetically reduced 3.0 Å resolutions. After reconstruction of the density for the entire protein, again using *phenix.maps*, the target density for the open state was extracted by masking the map about chain D using the *volutil* module of VMD.

**Figures 6** and **S8** suggests improvement in the refinement of CODH both at the high and intermediate resolutions. Similar to the ADK example, a higher number of replica improved the distribution of models across the range of CCs when fitting to a high resolution density map. But now, the best-fitted MDFF model improved from CC= 0.75 to 0.8 between *M_replicas_*=16 and 100, which an improvement of 6.7%, higher than the improvement of 3% seen in ADK over a similar range of replicas. In comparison to a single-replica MDFF of comparable length, where the final CC was 0.56, this improvement to 0.8 is even more dramatic, nearly 42%. Thus, the workflow on one hand scales with system size, while on the other benefits from the deployment of multiple replica simulations as the system size grows.

**Figure 6:**
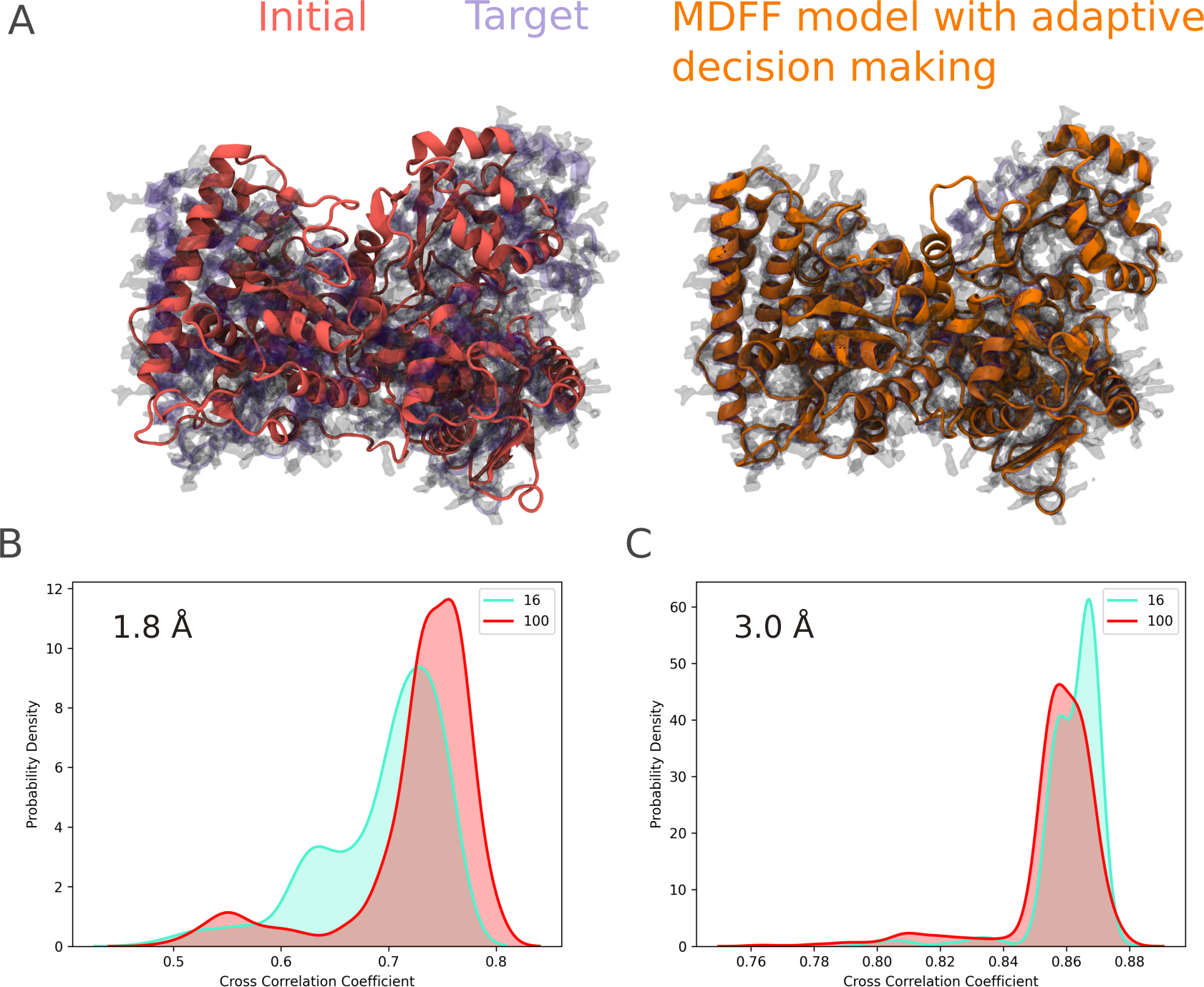
Robustness of R-MDFF protocol to system size. (A) Closed (red) to open (blue) are the initial and final (target) states of the dynamics for protein CODH. Flexibly fitting with R-MDFF to high resolution 1.8 Å EM density map. (B, C) Probability density of cross correlation coefficient for CODH, using the optimal parameters taken from ADK results at different resolutions - 1.8 Å (B) and 3.0 Å (C).

Our procedure of performing flexible fitting with on-the-fly adaptive decision making to transition from closed to open state, results in an ensemble within 2.0 Å RMSD to the experimentally determined “open” state comparing backbone atoms. This outcome is also comparable to our past refinement of CODH using a so-called cascade or simulated-annealing protocol, where the refined CODH model also reached within 2 - 2.5 Å of the open target. The larger number of replicas offer a search model greater number of opportunities to conform to density features in high resolution EM maps, and the *min* (1, *e*^Δ^*^ECC/KT^*) based selection-rule installed in R-MDFF enables avoiding of local minima in the CC space. Since the larger systems are prone to degeneracy of density features, we expect R-MDFF workflow enabled via EnTK to be more useful in overcoming the local minima and exhaustively sampling the conformational space as the size grows.

### 4.4 R-MDFF: Performance characterization

This section characterizes the computing performance of R-MDFF on HPC resources. We provide evidence that R-MDFF manages computing resources efficiently, with comparatively small overhead when running multiple replicas.

#### 4.4.1 Experiment Configuration

We designed 11 experiments to evaluate the efficiency of R-MDFF enabled via EnTK and we summarized their setup and results in Table 1. We utilized two biological systems—adenylate kinase (ADK) and carbon monoxide dehydrogenase (CODH)—running between 2 and 100 replicas (Rep), with varying simulation length (Sim. Len.) and resolution (Res.). Each experiment executed between 256 and 12800 tasks on PSC Bridges and ORNL Summit.

**Table 1:** Experiments to characterize R-MDFF performance. System: biological system name; Rep. (*M*): Total number of replicas between 2 and 100; Sim (ps): R-MDFF simulation length per iteration in picoseconds; Res. (Å): resolutions in Angstrom (high 1.8Å and intermediate 3.0Å); Resource: GPUs and CPU cores on OLCF Summit and CPU cores only on PSC Bridges2; Tasks: number of tasks for each experiment; OVH(s): Overhead of R-MDFF enabled via EnTK in seconds.

| Exp. ID | System | Rep. ($M$) | Sim. (ps) | Res. (Å) | Resource | Tasks | OVH (s) |
| --- | --- | --- | --- | --- | --- | --- | --- |
| 1 | ADK | 2 | 64 | 1.8 | Bridges (CPU) | 256 | $81.0 \pm 10$ |
| 2 | ADK | 4 | 32 | 1.8 | Bridges (CPU) | 512 | $126.0 \pm 10$ |
| 3 | ADK | 4 | 250 | 1.8 | Summit (GPU&CPU) | 512 | 92.0 |
| 4 | ADK | 8 | 160 | 1.8 | Summit (GPU&CPU) | 1024 | $105.27 \pm 18$ |
| 5 | ADK | 16 | 80 | 1.8 | Summit (GPU&CPU) | 2048 | $114.06 \pm 16$ |
| 6 | ADK | 32 | 40 | 1.8 | Summit (GPU&CPU) | 4096 | $109.33 \pm 10$ |
| 7 | ADK | 64 | 20 | 1.8 | Summit (GPU&CPU) | 8092 | $158.87 \pm 57$ |
| 8 | ADK | 100 | 10 | 1.8 | Summit (GPU&CPU) | 12800 | $266.98 \pm 245$ |
| 9 | CODH | 16 | 80 | 1.8 | Summit (GPU&CPU) | 2048 | $93.34 \pm 17$ |
| 10 | CODH | 16 | 80 | 3.0 | Summit (GPU&CPU) | 2048 | $99.44 \pm 20$ |
| 11 | CODH (long) | 100 | 80 | 1.8 | Summit (GPU&CPU) | 12800 | $113.61 \pm 20$ |

We characterized the performance of EnTK by measuring its overhead (OVH), i.e., the amount of time in which compute nodes are available but not used to execute tasks. Specifically, we separate between then time taken by the middleware (EnTK and its runtime system) to acquire resources, bootstrap the components and schedule the tasks; from the time taken by all the tasks to perform their scientific computation. Thus, OVH gives a simple but effective way to evaluate the cost of executing MDFF with EnTK and its components in terms of time spent to do everything else but science.

Experiment’s runs utilize up to 4 compute nodes on Bridges2 and 100, and execute each replica on a full compute node. On Bridges2, the NAMD MD engine uses 128 cores (AMD EPYC 7742 with of 256GB DDR4 memory), without GPU acceleration. Note that Bridges2 offers 24 compute nodes, each with 8 V100 GPUs accelerators but we decided to use only CPU resources due to their limited availability. On Summit, we run the CUDA-enabled NAMD MD engine on 6 NVIDIA V100 GPU accelerators per node. Different hardware platforms show wide performance gaps in time to solution but the cross-correlation is similar when using the same configurations.

We provide templates to allow users to replicate the experiments presented in this paper or as a starting point to create a run new experiments. The templates, written in YAML, store user-defined attributes for experiments and HPC resources separately, ensuring flexible analysis on diverse computing platforms. The source code and configuration parameters of the experiments are published on the R-MDFF Github repository. ^68^

#### 4.4.2 EnTK Overhead is Steady Across HPC Platforms

We measured the time spent by EnTK to bootstrap and clean up the execution environment. Those are overheads as they measure the time spent before and after the execution of the workflow’s tasks, when computing resources are already available. We measured the overheads on both Bridges and Summit, and at different scales.

Both bootstrap and clean up overheads are independent of the workflow scale as the time taken to manage the execution environment does not depend on the number of tasks executed in it. However, the bootstrap overhead can vary, depending on the filesystem performance and network latency, when serving packages and files during the bootstrapping process. We used a pre-configured environment to reduce the bootstrapping overhead by limiting the number of downloaded packages and the I/O operations required to build the execution environment of EnTK and the other RADICAL-Cybertools.

**Figure 7** shows that OVH is between 3% and 5% of the total execution time of the workflow presented in §3, across all our experiments. As summarized in Table 1, OVH is invariant of the number of replicas executed on Summit (150.90 *±* 115 seconds) and on Bridges (103.5 *±* 22.5 seconds) when running from 2 replicas to 100 replicas.

**Figure 7:**
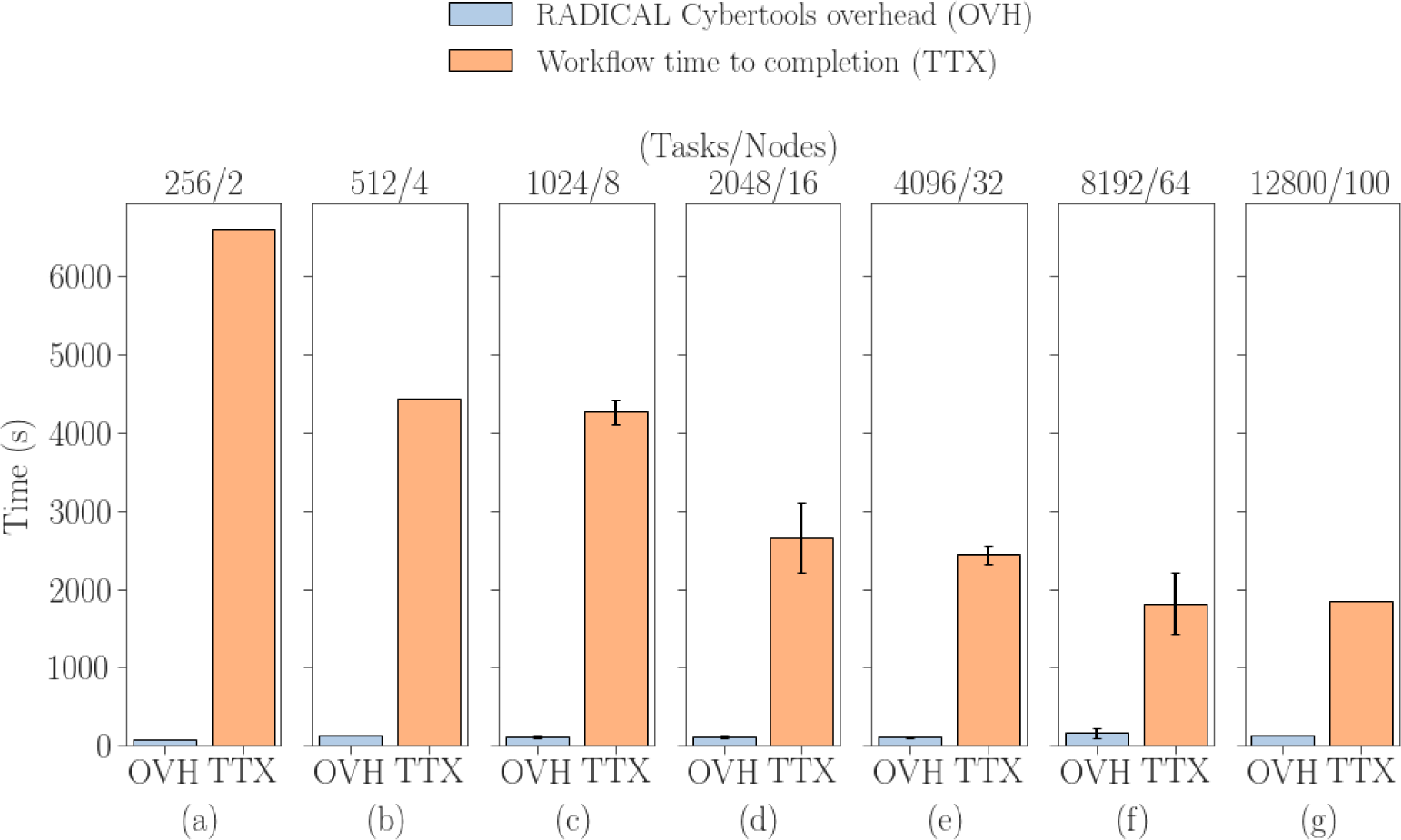
Performance of EnTK during flexible fitting with adaptive decision making. EnTK Overheads on PSC Bridges2 (a, b) and Summit (c - g) compute nodes. The total simulation time (equal to *t_steps/iteration_ × N_iteration_ × M_replicas_*) is 16ns. TTX tends to decrease with the increasing of the number of compute nodes.

Bridges2 shows three times larger overhead compared to Summit, mainly due to the different performance of the parallel filesystems: Lustre on Bridges2, GPFS on Summit. On Lustre, the initial access to files takes longer than continuous access because Lustre has to retrieve the location of the actual storage device over the network. The additional results from both platforms are reported in Supplementary Figures, **S9 and S10**.

### 4.5 Ensemble Refinement of pIX tetramer in Adenoviral Capsid of ChAdOx1

To demonstrate R-MDFF on an experimental cryo-EM density, the map of Oxford-AstraZenaca’s ChAdOx1 vaccine vector or capsid is re-refined, which we had originally published in ^81^ with PDB: 7RD1. We focus on the pIX tetramer subunit in particular. This subunit primarily provides structural stability to the icosahedral adenoviral capsid. ^82^ It also has additional functions in the viral life cycle. Conformational flexibility of the pIX tetramer compromises its local resolution, even when the overall resolution of the cryo-EM map is 3.07 Å. ^81^ By using Bayesian inference algorithms for resolution enhancement on RELION (version 3.1.1) ^83^ the density for the pIX tetrameric domain was already refined in our original submission (EMDB-24408). Now, to compare single and multi-model representations, we perform ensemble refinement of this homo-tetrameric protein. The exterior and interior region of the capsids asymmetric unit with the ensemble of pIX structures are illustrated in Figures 8 **A and B** respectively. We find that the collective CC of all members of the pIX ensemble considered simultaneously against the experimental density is twice that of the correlation derived from the distribution of MDFF models considered individually (Figure 8C). Hence, the lower resolution features of pIX is physically better determined using the multi-model description. The per-residue root mean square fluctuation (RMSF) was also computed from the refined ensemble (Figure 8D). The high fluctuations representative of the local disorder of pIX can only be captured with R-MDFF. A similar trend is seen in the SMOC scores. The regions of poor local resolution brings down the per-residue SMOC scores. ^84^ This trend is most evident for chain O between residues 10 to 30 (**Figure S11(A)**), where the local resolution as predicted by CryoRes ^85^ is the lowest (red region in **Figure S11(D)**). These TemPy2-SMOC scores ^84^ are calculated using a mean local resolution of 4 Å, which was estimated by CryoRes. ^85^ Single-model representation from traditional MDFF refinement is expected to offer a more skewed representation of the ensemble, ^3^ and hence underestimates the estimated fluctuations.

**Figure 8:**
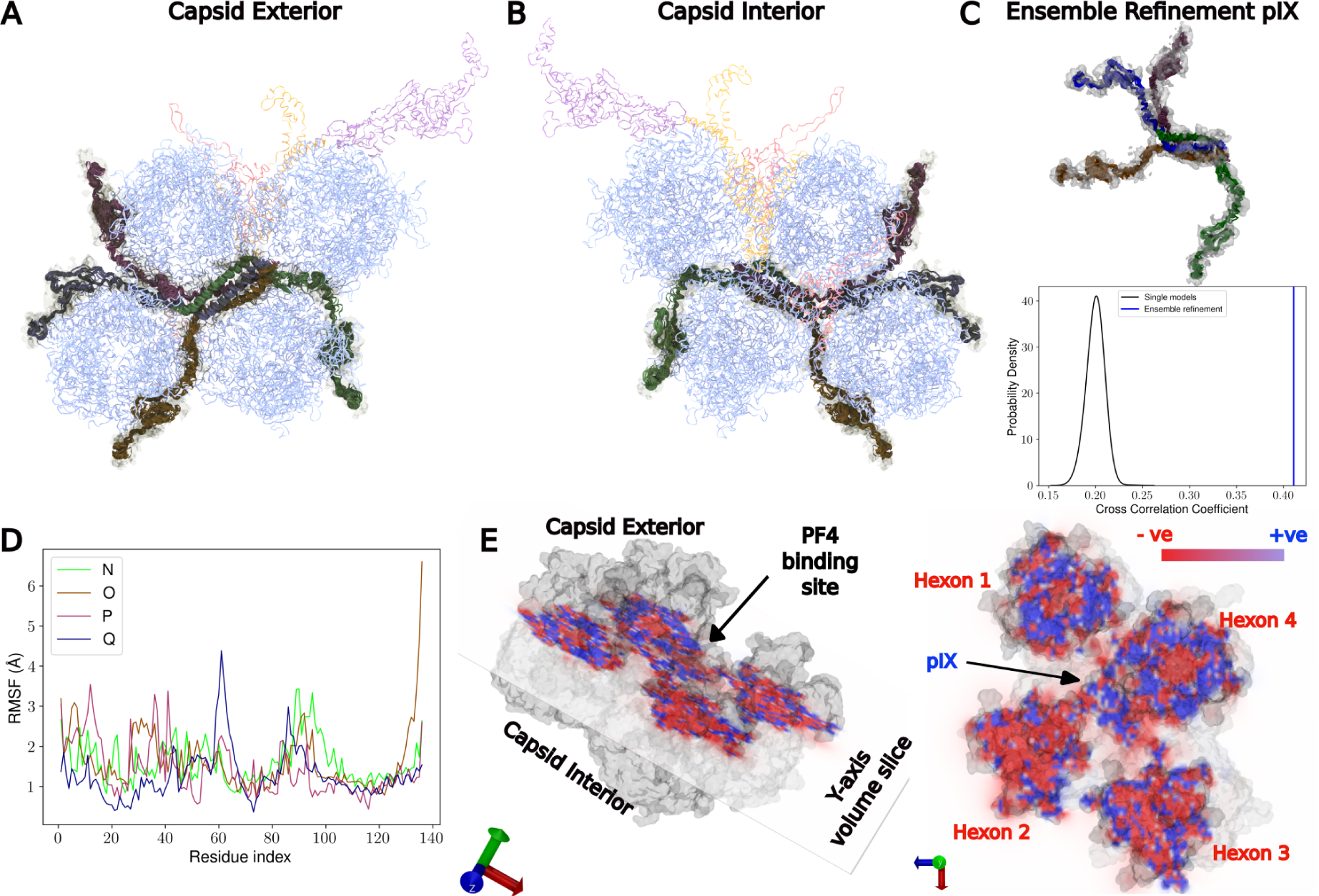
Ensemble refinement of pIX tetramer in ChAdOx1 asymmetric unit, using experimentally determined cryo-EM density. Asymmetric unit of ChAdOx1 where the different protein subunits are hexons (light blue), penton monomer (light pink), two copies of pVIII (light red), pIIIa (light orange) and result of ensemble refinement for tetrameric sub-unit pIX (green, dark pink, brown, navy blue) (A) capsid exterior (B) capsid interior. In (A, B) the EM density map for the pIX domain is show as a glass surface in tan. (C) Cross-correlation coefficient of pIX tetramer to the EM density is calculated for both single and multi-model representations. Here, multi-model is synonymous to the refined ensemble as shown by their trace, where each color corresponds to the different chains in pIX tetramer. The distribution of computed CC values between single models to the density is peaked at a lower value (black), when compared against the collection of all the single models fitted simultaneously (blue). This shows that an ensemble representation offers a more physically reliable model of the local density features than a single-model interpretation. The CC values are computed using the reported 3 Å resolution of the density map, which is certainly an overestimate, given poor local quality of the pIX region; this also reflects in alleviated values of the CC. Nonetheless, in the absence of a reliable estimate of the local resolution, we chose to use the reported global resolution of 7RD1. The CCs of both single and multiple-model fittings systematically increases by 1.5-fold when the resolution is set to 4 Å still maintaining a stronger correlation for the ensemble model. (D) Per-residue RMSF of individual chains in pIX tetramer indicate the local disorder in these flexible models. Analysis for RMSF per-residue is done using VMD-1.9.4, ^60^ enabled with python and numpy^87^ interface. (E) Average electrostatic potential isosurfaces (+5 *k_B_*T/e (blue) and -5 *k_B_*T/e (red)) for the hexon and pIX subunits in the ChAdOx asymmetric unit after ensemble refinement simulations. The hexon surfaces towards the capsid exterior are electronegative, while the pIX-rich region between the hexon trimers is electropositive. Electrostatic calculations are performed using APBS v3.0. ^88–90^ All illustrations for atomic model, pIX ensemble EM density and electrostatic potential maps are prepared using VMD-1.9.4. ^60^

Average electrostatic potentials of the resolved pIX conformations (Figure 8E) reveal a unique property of the ChAdOx1 capsid surface. The capsid consists of hexon proteins that are primarily negatively charged. They attract nonspecific binding with chemokine molecules, such as Platelet factor 4 (PF4) that has distinct positive and negative binding surfaces. This PF4-hexon binding is implicated in vaccine-induced thrombocytopenia of the ChAdOx1 vaccine. ^81^ Our resolution of pIX using the multi-model interpretation suggests that the PF4 binding of ChAdOx1 will be bimodal: the positive PF4 face will be focused away from the hexon-hexon interfaces, closer towards the core region of the subunits, and conversely, PF4s negatively charged apex will bind the pIX at the hexon interfacial regions, which is away from the hexon core. This asymmetry in binding mechanism arise from the ensemble interpretation of pIX.

## 5 Conclusions

Cryo-EM data of a protein represents an average of many two-dimensional images transformed to a three-dimensional density map. Classical methods in statistical mechanics such as MD fail to determine such an ensemble in finite length simulations, as structures remain trapped in deep potential wells corresponding to local dense points in the density maps. In view of the long-standing history of enhanced sampling and multi-replica free energy methods, to circumvent this algorithmic bottleneck of importance sampling and to decide the quality of an ensemble of protein structures on-the-fly, we present a framework for ensemble refinement of protein structures with adaptive decision making that improves both the quality of model and fit. We call this method R-MDFF. A refined protein ensemble offers, on one hand, the most probable structural representation based on available density information, while offering insights on protein conformational dynamics that are often ignored in traditional single-model interpretation derived from single-particle experiments.

The R-MDFF based workflow application allows adaptive decision-making for flexible fitting simulations by the the integration of correlation analysis with MD simulations. This workflow is implemented on two distinct heterogeneous high-performance national supercomputers facilities, Bridges2 and Summit. The workflow performs an user-defined number of iterative fitting and analysis tasks. This multi-replica scheme improves statistical significance, the quality of models over those derived from the traditional scheme of performing a single long MDFF simulation. Consequently, the new scheme arrives not just at the best-fit but a population of models with varied ranges of data-consistency. In addition, we show that R-MDFF enabled via EnTK, is well suited for heterogeneous extreme-scale high-performance computing environments ^86^ by managing resource utilization of GPU and CPU computing units and the workflow overhead for increased ensemble members. We also show that our approach would have a similar computational cost as the traditional single long MDFF simulation, but with a quick turnaround time (shorter wall time of workload), while exploring interesting regions in the density map. Larger system sizes that are more akin to cryo-EM structure determination offer further performance advantages. We continue to extend the capability of R-MDFF in complex applications in exascale high-performance computing environments.

## Data and Software Availability

The data including input scripts, parameter files, topology files, and trajectories are available from the corresponding author upon request. The code for the R-MDFF workflow is made publicly available on GitHub (https://github.com/radical-collaboration/MDFF-EnTK) and the data for this study is made publicly available on Zenodo (https://doi.org/10.5281/zenodo.7695738). The instructions to download and install the source code for R-MDFF is publicly available on GitHub. Kindly refer https://github.com/radical-collaboration/MDFF-EnTK/blob/master/install.md for further details.

## Supporting Information Available

The Supporting Information (SI) includes a trace plot for the cross correlation coefficient, calculated after every iteration in the R-MDFF protocol to fit ADK in high, intermediate and low resolution electron denisty maps. The SI also includes MolProbity scores for models from R-MDFF trajectories and the performance of the R-MDFF algorithm when applied to larger biomolecular systems such as CODH.

## Acknowledgement

A.S. acknowledges start-up funds from SMS and CASD at Arizona State University, CAREER award from NSF (MCB-1942763), and an NDEP grant from the Department of Defense. This work used the Extreme Science and Engineering Discovery Environment (XSEDE), which is supported by National Science Foundation grant number ACI-1548562. The RADICAL Lab acknowledges NSF Awards 1835449 and 1931512. The benchmarks were also carried out using the resources of the OLCF at Oak Ridge National Laboratory, which is supported by the Office of Science at DOE under Contract No. DE-AC05-00OR22725, made available via the INCITE program. J.W.V acknowledges the support from the National Science Foundation Graduate Research Fellowship under grant number 2020298734. D.S. acknowledges the important discussions and feedback from Dr. Alexander T. Baker (Accession Therapeutics) and Dr. Chun Kit Chan (Arizona State University) for the structural modeling, refinement and APBS calculations for the pIX protein in ChAdOX1.

## TOC Graphic

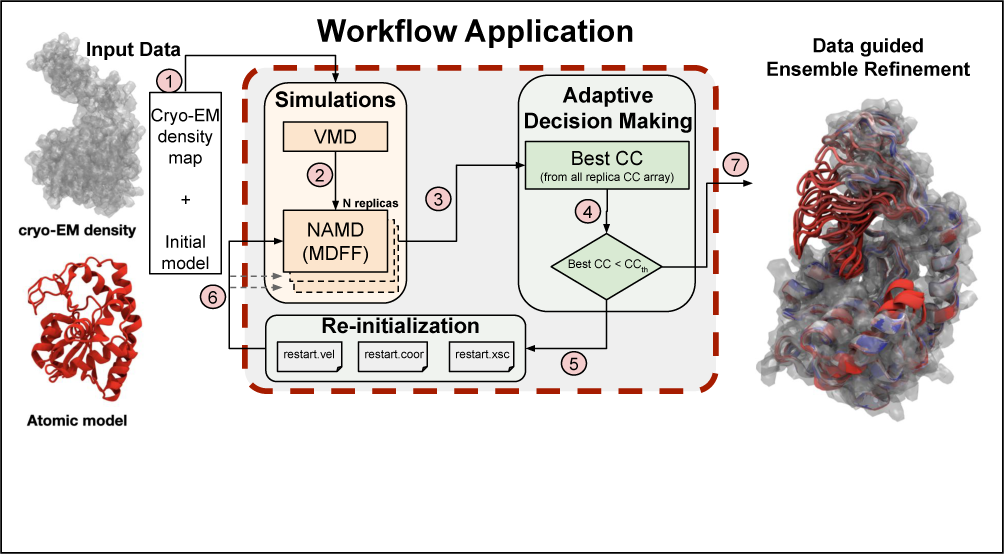

## Notes

### Competing Interest Statement

The authors have declared no competing interest.

### Summary of Updates

New data and analysis included in main text and supplementary information.

