## Supplemental Information for "Adaptive Ensemble Refinement of Protein Structures in High Resolution Electron Microscopy Density Maps with Radical Augmented Molecular Dynamics Flexible Fitting"

### Comparing cross correlation coefficient after R-MDFF for flexible fitting ADK in different resolution electron density maps

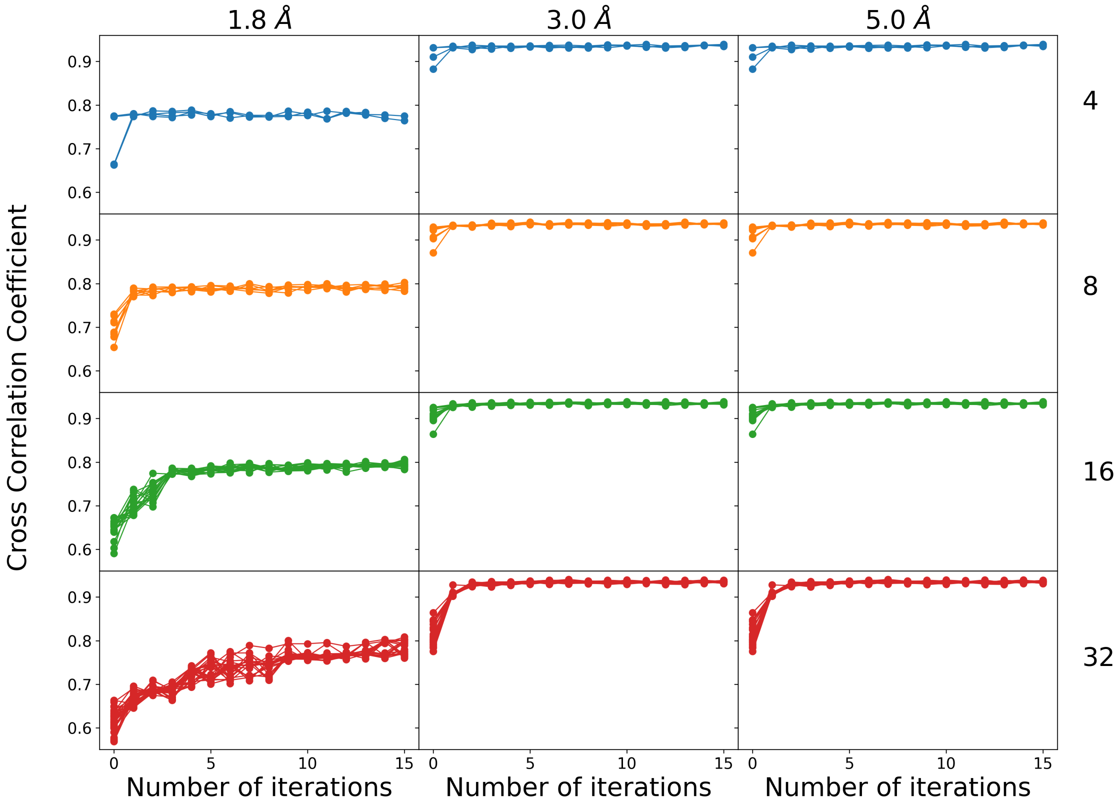


Figure S1: **Cross correlation coefficient every iteration during flexibly fitting ADK with R-MDFF at different resolution density maps.** Data presented for high resolution (1.8 Å), intermediate resolution (3.0 Å) and low resolution (5.0 Å) cryo-electron microscopy density maps, for different ensemble members, 4 (blue), 8 (orange), 16 (green) and 32 (red) respectively. We find that replica number directly correlates with we have larger variabilty amongst ensemble members for high resolution density map.

### Comparing RMSD between structures from R-MDFF refinement using 400 replicas to the reference model of ADK fitted in density from 16-replica dataset.

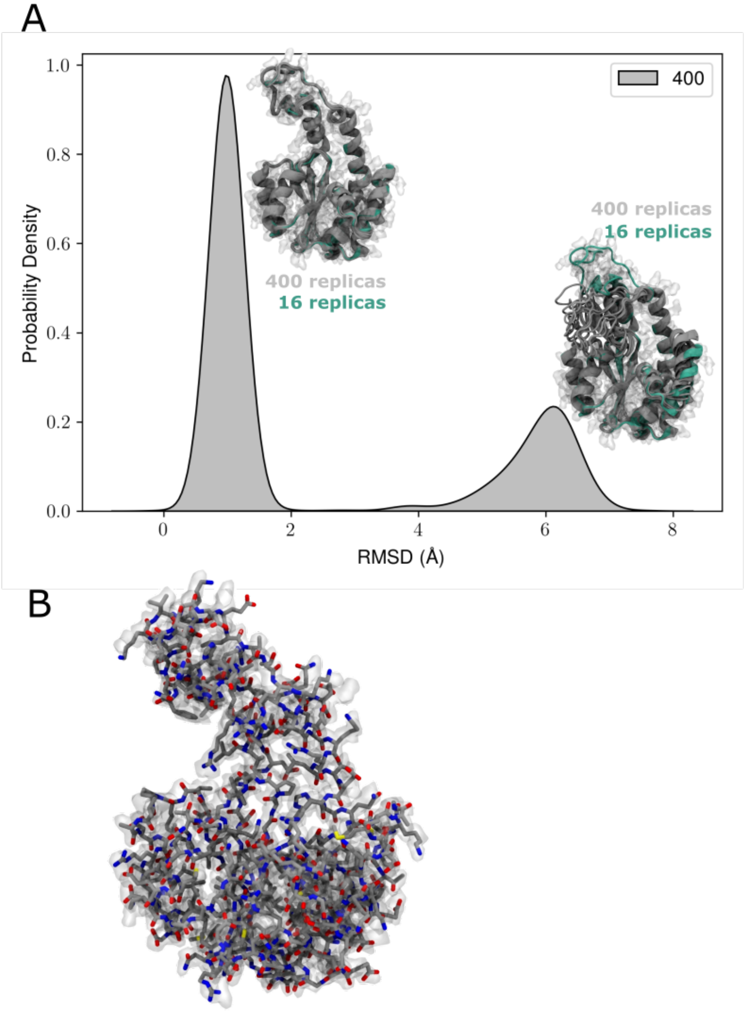


Figure S2: **RMSD comparison between structures from 400 replica R-MDFF simulation to fitted structure from 16 replica model.** (A) Bimodal probability distribution for RMSD calculated between ADK structures obtained after R-MDFF refinement using 400 replicas (gray) to a fitted model from 16 replica ensemble (aqua green). The RMSD calculation is performed using sidechain atoms with hydrogen in protein ADK. (B) Illustration of a structure from the high CC (Fig. 5A) population for ADK after flexible fitting using R-MDFF with 400 replicas with synthetic density of 1.8 Å. As expected, the RMSD around 1 Å indicates that high CC population shown in Fig. 5A, are similar to the ensemble members in the case of 16 replicas.

### MolProbity calculation after flexibly fitting ADK to high resolution density map

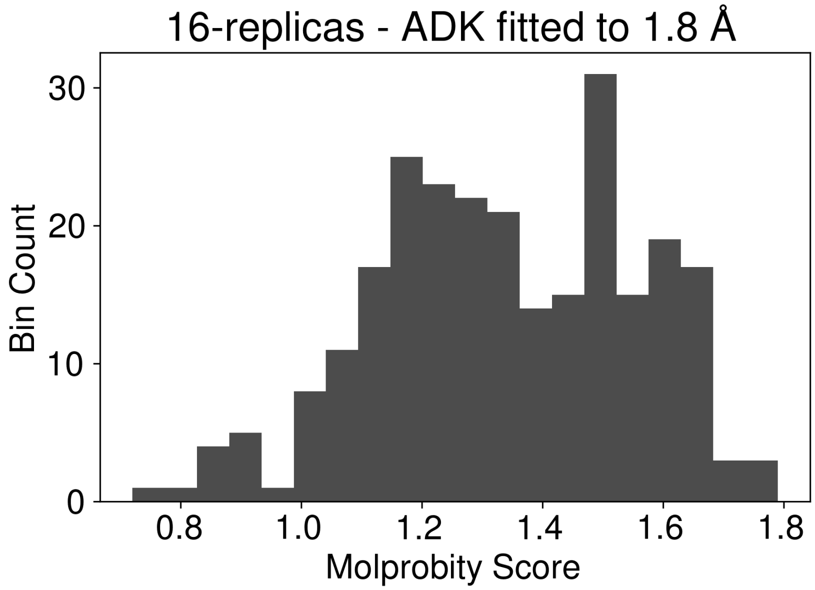


Figure S3: Overall MolProbity scores for 16 replicas of ADK flexibly fitted to 1.8 Ådensity map. Scores are calculated using Phenix. molprobity package.[S1]

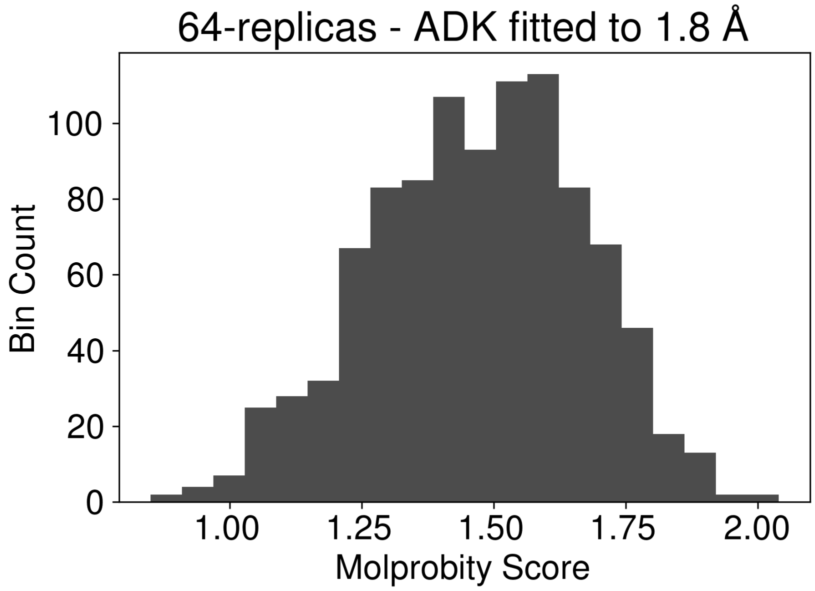


Figure S4: Overall MolProbity scores for 64 replicas of ADK flexibly fitted to 1.8 Ådensity map. Scores are calculated using Phenix. molprobity package.[S1]

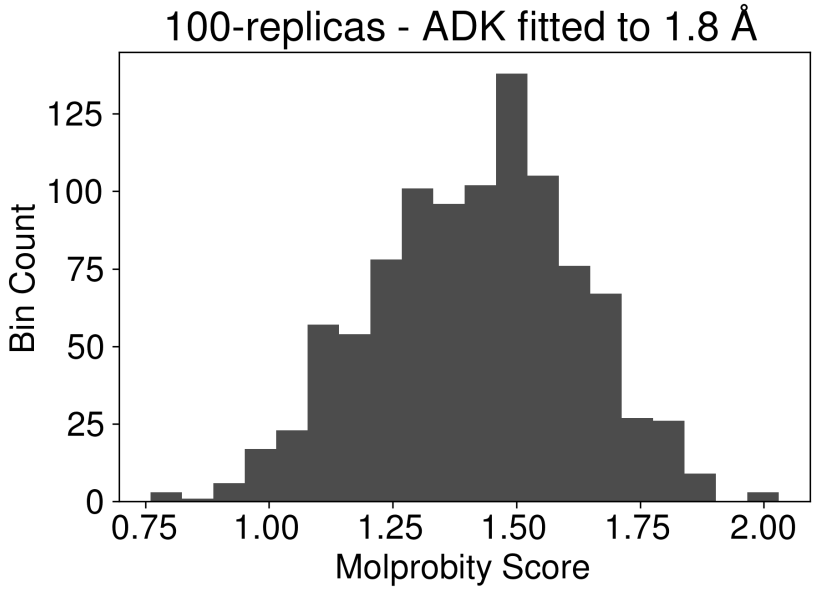


Figure S5: Overall MolProbity scores for 100 replicas of ADK flexibly fitted to 1.8 Ådensity map. Scores are calculated using Phenix. molprobity package.[S1]

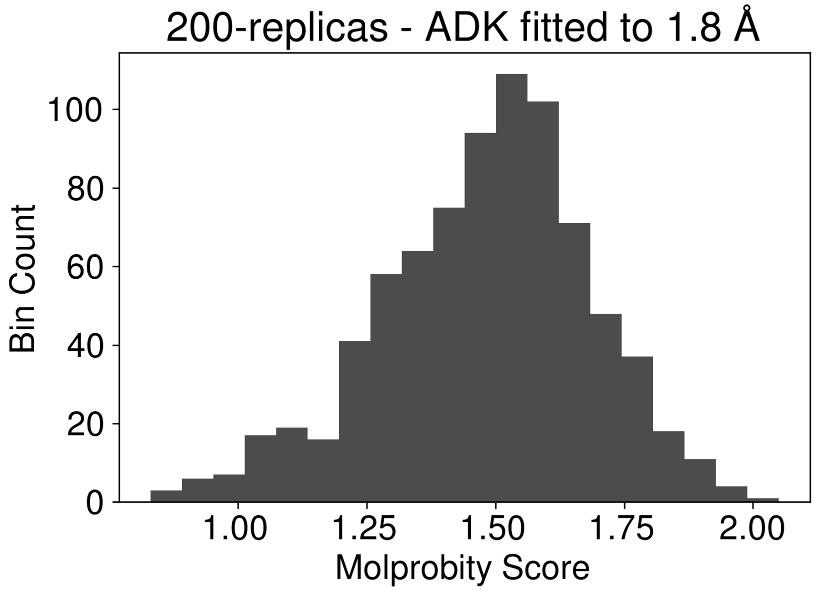


Figure S6: Overall MolProbity scores for 200 replicas of ADK flexibly fitted to 1.8 Ådensity map. Scores are calculated using Phenix. molprobity package.[S1]

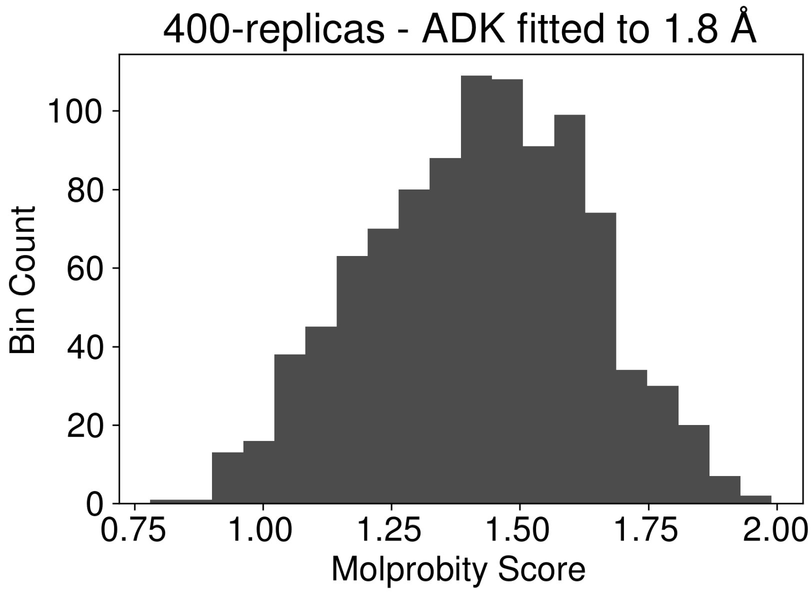


Figure S7: Overall MolProbity scores for 16 replicas of ADK flexibly fitted to 1.8 Ådensity map. Scores are calculated using Phenix. molprobity package.[S1]

### R-MDFF results for carbon monoxide dehydrogenase (CODH) using 100 replicas

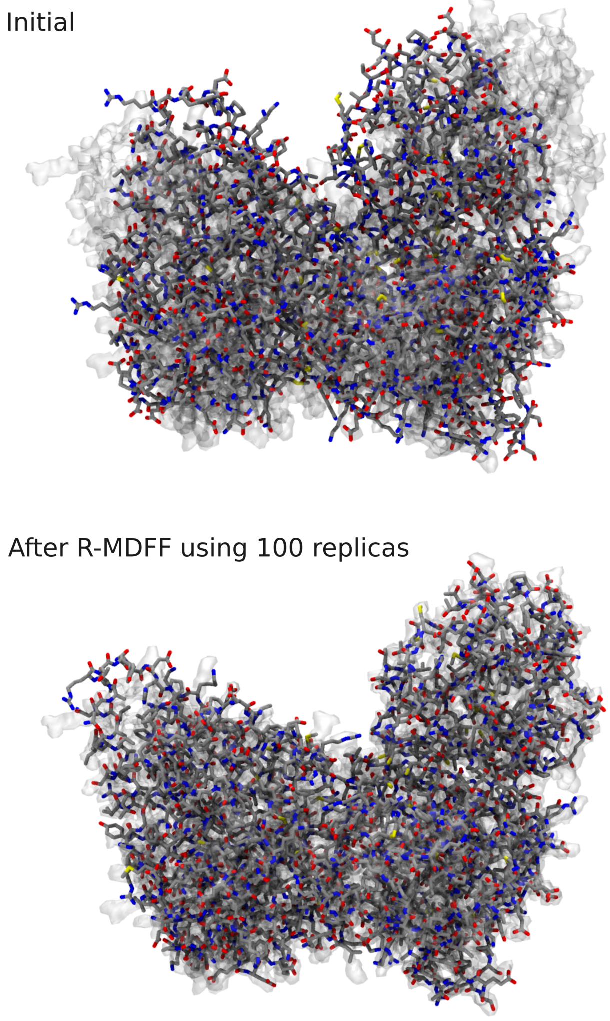


Figure S8: **Illustrating R-MDFF results after CODH is fitted to a 1.8 Å density map using 100 replicas**. The initial and after R-MDFF refinement. CODH structure is shown using the Licorice representation in VMD, to clearly illustrate the side chains before and after flexible fitting. The high resolution density map is shown as a transparent surface for clarity.

### Performance of R-MDFF for CODH on HPC clusters

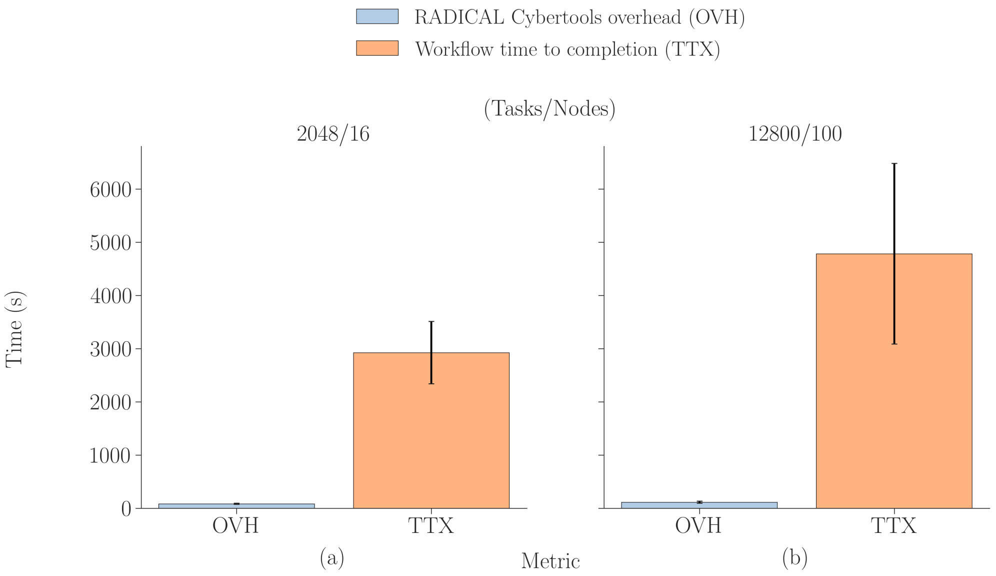


Figure S9: RADICAL Overheads for CODH on 16 and 100 Summit compute nodes. The simulation time per iteration is 80 ps and TTX tends to decrease with the increasing of the number of compute nodes.

Figure S9 shows the RADICAL overheads of running CODH with 16 replicas and 100 replicas in which $99.44 \pm 20$ seconds and $113.61 \pm 20$ seconds are measured respectively. The bar plots with a light blue color (OVH label on X-axis) indicate the overhead in seconds of completing the R-MDFF workflow, and the bars with orange color (TTX label on X-axis) report estimated workflow time to completion for sixteen iterations. The experiments id are 7, 8 and 9 in Table 1, which corresponds to Figures S9 and S10.

Figure S10 shows the RADICAL overheads of running 2 and 4 replicas on PSC Bridges in which $636.18 \pm 10$ seconds and $332.39 \pm 10$ seconds are measured respectively. These overheads include 87 seconds and 42 seconds startup time (bootstrap) for 2 and 4 replicas and we believe that initial access delay performance on Lustre file system is fluctuated.

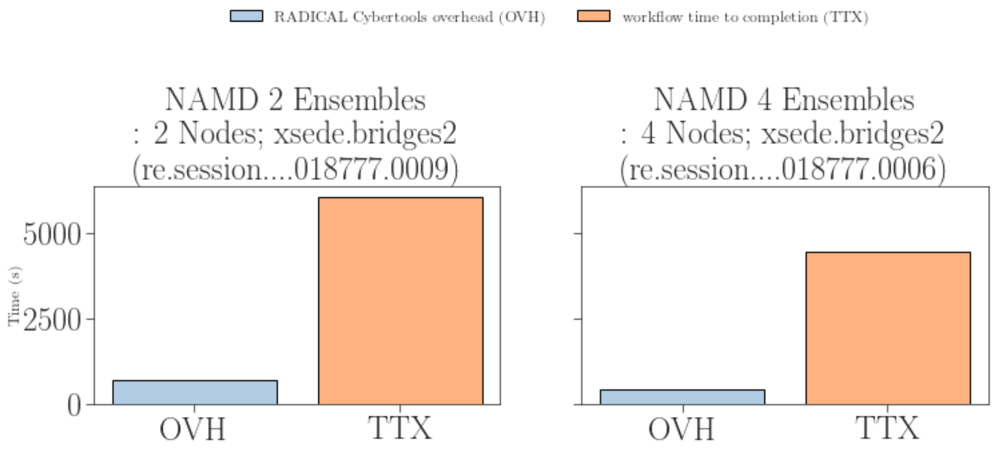


(a) (b)

Figure S10: RADICAL Overheads on PSC Bridges2 compute nodes. (ADK, 1.8A, 2/4 replicas, 2048 ps timescale)

### Local cross correlation calculation and SMOC score analysis for pIX refinement.

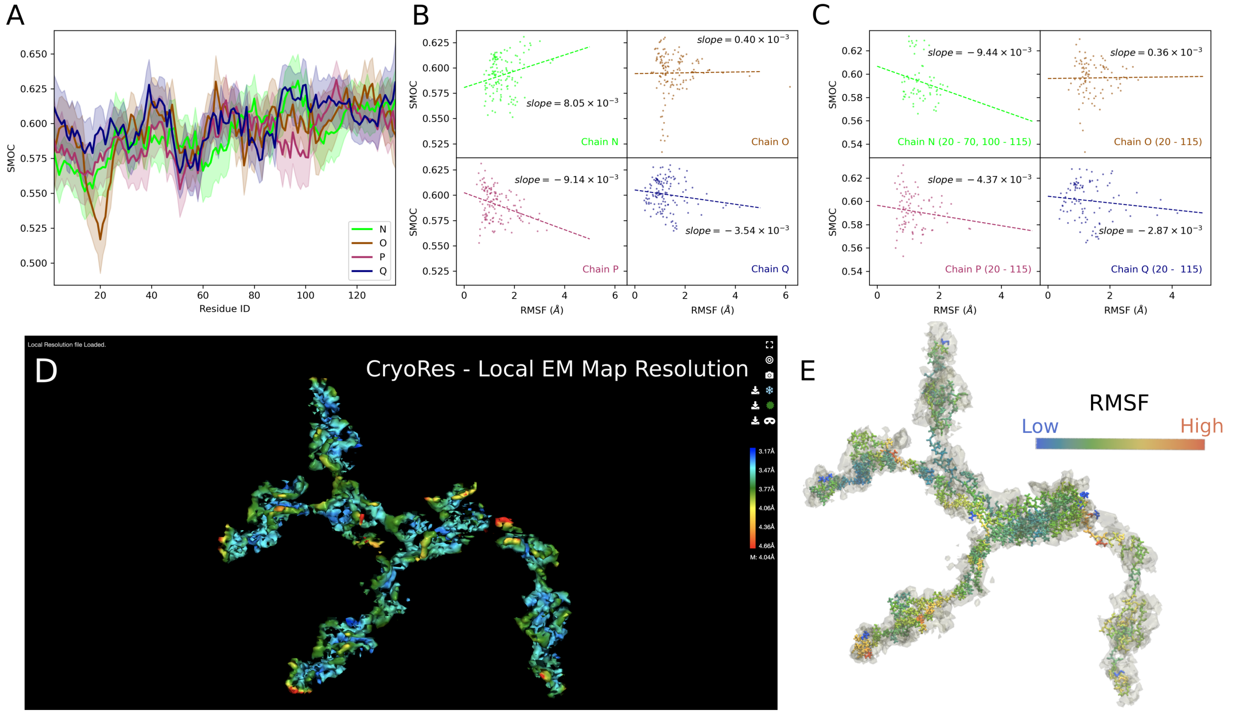

Figure S11: **Local cross correlation coefficient and residue based SMOC score of pIX ensemble.** (A) Using the density map for the pIX region in ChAdOx1[S2] asymmetric unit (Fig. 8 A - C), the per-residue SMOC[S3] score is calculated for the entire ensemble. The different chains in the tetramer pIX sub-unit are N (green), O (brown), P (dark magenta), Q (dark blue). The mean SMOC score for the ensemble is shown in thick solid lines, while the standard deviation is illustrated by a transparent region. The SMOC score indicates the local goodness of fit of the model with respect to the underlying density map and higher numbers indicate a better fit. (B) Correlation between per-residue SMOC and RMSF calculations - where all residues in individual chains are considered in linear regression and (C) performing linear regression[S4] ignoring the N and C-terminal regions for all chains. In addition for chain N, specifically residues in the "discontinuous" density region are removed. For chains P and Q, we observe a negative slope between per-residue SMOC and RMSF scores. Generally, higher fluctuation of the local regions in the protein correspond to the local low resolution in the density map. Expectedly, the correlation between local SMOC and RMSF trend is weak. This is because of the limited number of MD models used for the RMSF computation, which makes the RMSF estimates noisy. Also, the map-model correlation changes on a different, potentially slower temporal scale than real atomistic fluctuations.[S5] (D) CryoRes[S6] local resolution estimation using a deep learning based algorithm. On the relative scale, blue indicates high resolution, while red indicates low resolution. (E) Atomic structure for the last frame of the ensemble refinement trajectory on individual chains in pIX colored by their per-residue RMSF (calculated in Fig. 8D). Here, blue indicates low and red refers to high values of fluctuation in the structure. The deep learning based local resolution prediction correlates well with the fluctuations observed in the ensemble during refinement.

(S4) Virtanen, P.; Gommers, R.; Oliphant, T. E.; Haberland, M.; Reddy, T.; Cournapeau, D.; Burovski, E.; Peterson, P.; Weckesser, W.; Bright, J.; van der Walt, S. J.; Brett, M.; Wilson, J.; Millman, K. J.; Mayorov, N.; Nelson, A. R. J.; Jones, E.; Kern, R.; Larson, E.; Carey, C. J.; Polat, İ.; Feng, Y.; Moore, E. W.; VanderPlas, J.; Laxalde, D.; Perktold, J.; Cimrman, R.; Henriksen, I.; Quintero, E. A.; Harris, C. R.; Archibald, A. M.; Ribeiro, A. H.; Pedregosa, F.; van Mulbregt, P.; SciPy 1.0 Contributors, SciPy 1.0: Fundamental Algorithms for Scientific Computing in Python. *Nature Methods* **2020**, *17*, 261–272, DOI: 10.1038/s41592-019-0686-2.

(S5) Vant, J. W.; Sarkar, D.; Streitwieser, E.; Fiorin, G.; Skeel, R.; Vermaas, J. V.; Singharoy, A. Data-guided Multi-Map variables for ensemble refinement of molecular movies. *Journal of Chemical Physics* **2020**, *153*, DOI: 10.1063/5.0022433.

(S6) Dai, M.; Dong, Z.; Xu, K.; Zhang, Q. C. CryoRes: Local Resolution Estimation of Cryo-EM Density Maps by Deep Learning. *Journal of Molecular Biology* **2023**, *435*, 168059, DOI: 10.1016/j.jmb.2023.168059.
